# Landscapes of missense variant impact for human superoxide dismutase 1

**DOI:** 10.1101/2025.02.25.640191

**Authors:** Anna Axakova, Megan Ding, Atina G. Cote, Radha Subramaniam, Vignesh Senguttuvan, Haotian Zhang, Jochen Weile, Samuel V. Douville, Marinella Gebbia, Ammar Al-Chalabi, Alexander Wahl, Jason Reuter, Jessica Hurt, Adele Mitchell, Stephanie Fradette, Peter M. Andersen, Warren van Loggerenberg, Frederick P. Roth

## Abstract

Amyotrophic lateral sclerosis (ALS) is a progressive motor neuron disease for which important subtypes are caused by variation in the Superoxide Dismutase 1 gene *SOD1*. Diagnosis based on *SOD1* sequencing can not only be definitive but also indicate specific therapies available for *SOD1*-associated ALS (SOD1-ALS). Unfortunately, SOD1-ALS diagnosis is limited by the fact that a substantial fraction (currently 26%) of ClinVar SOD1 missense variants are classified as “variants of uncertain significance” (VUS). Although functional assays can provide strong evidence for clinical variant interpretation, SOD1 assay validation is challenging, given the current incomplete and controversial understanding of SOD1-ALS disease mechanism. Using saturation mutagenesis and multiplexed cell-based assays, we measured the functional impact of over two thousand SOD1 amino acid substitutions on both enzymatic function and protein abundance. The resulting ‘missense variant effect maps’ not only reflect prior biochemical knowledge of SOD1 but also provide sequence-structure-function insights. Importantly, our variant abundance assay can discriminate pathogenic missense variation and provides new evidence for 41% of missense variants that had been previously reported as VUS, offering the potential to identify additional patients who would benefit from therapy approved for SOD1-ALS.

## Introduction

Amyotrophic lateral sclerosis (ALS; MIM: 105400) is a heritable mainly-autosomal-dominant neurodegenerative disease with a lifetime prevalence of 1 per 300 people^1–3^. *SOD1* encodes the ubiquitously expressed free-radical scavenging enzyme Cu^2+^/Zn^2+^ superoxide dismutase (MIM: 147450 and EC: 1.15.1.1), variants of which appear in 9-23% of diagnosed familial ALS (fALS) cases, and at least 2- 5% of diagnosed sporadic ALS (sALS) cases depending on the population^4–7^.

Historically, ALS diagnosis is delayed for an average of one year after symptom onset^8,9^. However, genetic analysis now offers the potential for earlier diagnosis of ALS or carrier status and can identify subsets of patients who may benefit from specific therapeutic interventions^10,11^. Classifying the pathogenicity of rare missense variants remains challenging. Of the 156 missense variants of SOD1 that have been reported in ClinVar, 26% have been categorized as ‘variants of uncertain significance’ (VUS) and none have been interpreted as likely benign or benign^12^. More broadly, for variants across many disease genes, validated functional assays can provide strong evidentiary value under current American College of Medical Genetics and Genomics and Association for Molecular Pathology (ACMG/AMP) guidelines^13^. Cell-based functional assays, either in cultured human cells or more tractable eukaryotic model systems, like yeast, can be used to detect variant functional impacts^14^.

In any case, small-scale functional assays are resource-intensive and typically ‘‘reactive,’’ performed only after (and often years after) the first clinical presentation of a variant. In contrast, computational methods can predict the impact of all missense variants ‘‘proactively,’’ in advance of the first clinical presentation. While computational predictors are steadily improving, these are better suited to detect pathogenic missense variants that cause loss of function^15^. Currently, there are no published variant interpretation guidelines for *SOD1*, and computational methods, as outlined in the ACMG/AMP recommendations for variant interpretation^13^, provide at best supporting evidence. Although recent studies have suggested that greater weight should be afforded to computational predictors of pathogenicity^16,17^ this has not been validated for *SOD1* in particular.

Within the past decade, it has become possible to proactively test missense variants at large scale using massively multiplexed cell-based assays, yielding exhaustive sequence-function maps (‘variant effect maps’) that describe functional impacts of thousands of variants^18^. These maps can accurately identify functional variants, even for variants that had not yet been clinically observed before the map was produced. Moreover, functional evidence from variant effect maps has already begun to assist variant reclassification^18,19^.

Here we employ highly-multiplexed cell-based assays to proactively and systematically measure all missense variant impacts for the human SOD1 protein using multiplexed assays in both yeast and human cells. We find that the resulting impact scores of nearly all possible missense variants correspond well with prior knowledge about the SOD1 protein structure and with known patterns of mutational tolerance, and also point to additional sequence-structure-function relationships. Finally, we demonstrate that variant effect map scores provide new large-scale evidence to classify the pathogenicity of *SOD1* alleles.

## Results

### 2.1 Implementing scalable functional assays for SOD1 missense variants

For *SOD1*, previous small-scale cell-based assays have been based on the premise that SOD1 variant pathogenesis is via toxic gain of function variants^20–22^. More specifically, the model suggests that toxic gain of function, whether from SOD1 variation or interaction with other proteins such as FUS and TDP-43, leads to the polymerization of SOD1^23^. Although cell-based aggregation assays have been used as evidence in support of clinical variant annotation, these assays have either used patient cells (which is not scalable), or required either transient transfection^22^ or lentiviral integration^20^, yielding SOD1 variant expression that potentially goes well beyond physiological levels. We therefore sought to identify a scalable cell-based assay for the propensity of SOD1 variants to aggregate which employed moderate SOD1 expression levels. Unfortunately, despite pursuing both yeast and human cell based assays under a variety of sensitizing conditions we were unable to develop such an assay (see Document S1).

Although many pathogenic SOD1 variants associated with ALS exhibit reduced function, variants that completely ablate SOD1 enzymatic activity have been linked to a progressive, adult-onset ALS phenotype^24^, an infantile SOD1 deficiency syndrome (ALS-phenotype)^25^ and aneurological disorder distinct from ALS^26–28^. Pathogenic variants of SOD1 have been described as often having both gain of function (aggregation) effects and partial loss of enzymatic activity. A complicating factor is that ALS-associated variants can display varied expression levels and decreased half-lives^6,26,29–31^. Moreover, both expression and enzymatic activity of SOD1 variants can be tissue-dependent^32^. We therefore next sought to assess the impact of SOD1 variants on both enzymatic function and protein abundance.

First, to investigate the functional impact of missense variation in SOD1 at scale, we implemented a previously-validated humanized yeast model. Yeast *sod1Δ* strains are known to have growth defects under heat-stress (38°C) growth conditions, and we confirmed that expression of human *SOD1* rescues this growth defect^33,34^. We evaluated this assay by measuring growth rescue for a small set of high-confidence pathogenic variants as defined by multiple ClinVar submitters (Figure S1). This set of variants included some known to exhibit both aggregation and loss-of-enzymatic activity effects (p.His47Arg and p.Asn132Lys), some known to exhibit aggregation for which the impact on activity had not been previously reported (p.Gly94Arg, p.Ile114Thr and p.Leu145Phe), and used both p.Asn20Ser and p.Gly130Ser as a proxy-benign control (given the absence of any annotated benign variants in ClinVar) drawn from gnomAD. Each variant, together with a positive control WT human *SOD1* and an empty vector control, were expressed in the yeast *sod1Δ* strain under thermal stress. We observed complementation for all non-pathogenic variants (100% precision) while observing lack of complementation for two of five pathogenic variants (40% recall). Furthermore, no SOD1 variants showed toxicity under normal growth conditions, which was unexpected for the subset of variants (p.His47Arg, p.Gly94Arg, and p.Ile114Thr) with known gain of function effects.

Next, to assess the impact of SOD1 variants on protein abundance, we implemented a Variant Abundance by Multiplexed Parallel Sequencing assay (VAMP-Seq) assay, in which the protein of interest is fused to GFP. Using this approach, variants causing misfolding or destabilization followed by degradation can be detected on the basis of reduced GFP expression^35^. Constructs containing *SOD1*-*GFP* variants were introduced into human HEK293T cells and integrated at a ‘landing pad’ containing a *Bxb1* recombination site flanked by an inducible doxycycline-inducible promoter on one side of the *Bxb1* site, enabling expression of a BFP gene on the other side. As a result, cells with integration events (which can be enriched by sorting for BFP- cells) each express one copy of a single variant SOD1-GFP construct^35^. Because SOD1- ALS is a dominant disease that generally presents in patients who are heterozygous carriers of a pathogenic variant, the endogenous copy of *SOD1* was not knocked out.

We initially subjected a small set of high-confidence (multiple submitters on ClinVar) reported pathogenic gain of function (aggregating) SOD1 variants to this assay, measuring GFP abundance in each integrated cell line via flow cytometry (Figure S2). This set of variants included some known to exhibit both aggregation and loss-of-enzymatic activity effects (p.Ala5Val, p.His47Ala, p.Gly86Arg, p.Gly94Ala, p.Arg116Gly, p.Leu145Phe^36–38)^. We included p.Gly130Ser as a proxy-benign control drawn from gnomAD. All variants we tested showed distributions of GFP intensity that were lower than that of WT SOD1, although the pathogenic p.His47Arg and p.Leu145Phe variants, and Gly130Ser proxy-benign control showed intermediate GFP levels (higher than all pathogenic variants but lower than WT).

### 2.2 Systematically testing SOD1 variant effects on enzymatic activity and protein abundance

To measure functional impacts for all possible missense SOD1 variants, we next adapted the above-described total enzymatic activity and abundance assays as multiplexed assays of variant effects, using the previously-described TileSeq framework (Figure 1A)^39^. As an initial step, we constructed mutagenized libraries of SOD1 variants (with and without C-terminally fused GFP) by using a large-scale oligo-directed codon mutagenesis protocol (see Methods). Each mutagenized library was initially generated as a pool of amplicons, and transferred *en masse* via two steps of recombinational subcloning (see Methods) into one of two alternative ‘destination’ vectors. Care was taken in this process to maintain pool complexity such that the average variant appears in at least 50 independent clones. Mutagenized destination vector libraries were introduced into the appropriate yeast or human assay cells, and subjected to the appropriate selection procedure:

**Figure 1:**
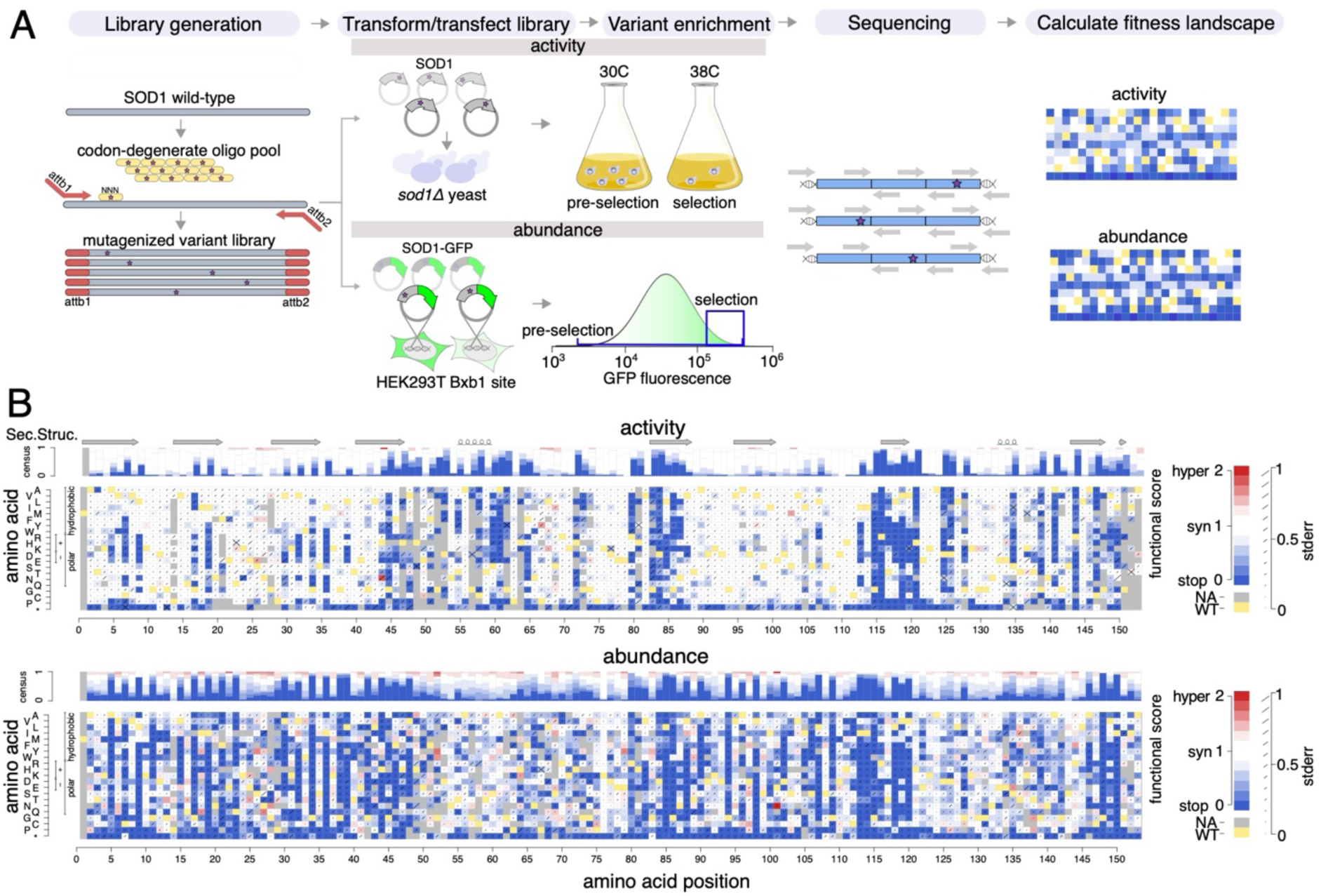
Generation of SOD1 missense variant effect maps. (A) Overview of process to generate variant effect maps. (B) SOD1 variant effect maps measuring total enzymatic activity (top) and abundance (bottom). Box color either indicates the WT residue (yellow); a substitution with damaging (blue), tolerated (white), or above-WT (“hyper”; red) functional score; or missing data (gray). Consensus tracks summarize the map scores by position. β-strands are indicated by arrows, and alpha helices by loops.

First, to assess the loss-of-function effects of SOD1 variants in parallel, the mutagenized library without fused GFP was cloned *en masse* into yeast expression vectors (designed to express SOD1 using the moderate-strength yeast *ADH1* promoter) and transformed into the yeast *sod1*Δ strain. In order to maintain the expression plasmid, the transformed yeast strain pool expressing human SOD1 variants was maintained in Leu-synthetic media. To enrich for SOD1 variants that retain enzymatic function, cells were grown competitively at the restrictive (38°C) temperature at which SOD1 activity is essential. The relative frequencies of variants were measured in cell pools both pre-selection and post-selection conditions.

Second, to assess SOD1 variant impacts on protein abundance, we used the VAMP-seq method in cultured human cells to measure the steady-state abundance of human SOD1 variants. The mutagenized library was designed such that each SOD1 variant clone has a C-terminal GFP fusion followed by an internal ribosomal entry site and mCherry (as a marker of both transfection and construct expression level). This library was recombined into HEK293T cells containing the *Bxb1* landing pad, simultaneously disrupting BFP expression at the landing pad and enabling expression of SOD1-GFP and mCherry via the TetOn promoter upstream of the Bxb1 site. Stable recombinant integrant cells harboring SOD1 variants (BFP-, mCherry+ cells) were then isolated by FACS and expanded. To enrich for SOD1 variants that retain protein abundance, we selected for a population with both high GFP and mCherry expression (“post-selection” condition), and measured the relative frequencies of variants in these cells relative to all SOD1 integrant cells (the “pre-selection” condition).

For both total enzymatic activity and abundance assays, we obtained functional scores of each SOD1 variant by measuring the relative abundance of that variant in the ‘post-selection’ relative to the ‘pre-selection’ conditions, using the previously described TileSeq approach^40^. Briefly, five ∼150bp tiles, collectively covering the entire gene coding region, were subjected to “duplex sequencing” (in which each single-molecule-derived ‘colony’ of PCR products is sequenced on both strands), allowing lower base-calling error. Each nucleotide position had a sequencing depth of at least 1.3 million reads, and we considered variants seen at a frequency of at least 10 counts per million reads in the pre-selection condition to have been well-measured. For both assays, this criterion was met by 96% of all possible missense variants and 99% of those amino acid substitutions that are achievable via a single-nucleotide variant (SNV) (Figure S4A). The average number of *SOD1* codon substitutions per cell was estimated at 0.64 for activity and 0.83 for the abundance map (Figure S4B). The uncertainty (standard error) for each functional score was also estimated as previously described^41,42^, based on trends in the extent of agreement between biological replicates as a function of variant frequency in the pre-selection condition. After the filtering described above, the scores for all variants were rescaled such that 0 corresponds to the median of nonsense variants and 1 to the median of synonymous variants.

Biological replicate scores showed high agreement for both the total enzymatic activity (Pearson’s *R*=0.81) and abundance (Pearson’s *R*=0.79) maps (Figure S4C). Among the 2605 and 2632 missense variants measured in each map, respectively, we observed the score distribution for nonsense variants to be strongly shifted downward relative to synonymous variants (Figure S4D). To generate the final scores, we averaged the biological replicates for each variant and estimated error in the final score as described in Methods. A bimodal distribution of missense variants was observed for both maps, with modes that fell close to those of nonsense and synonymous variants, presumably corresponding to either deleterious or neutral effects, respectively (Figure S4D). The peaks corresponding to deleterious variants in the activity and abundance maps contained ∼14% and ∼18% of missense variants, respectively. Figure 1B visualizes the complete variant effect maps for both total enzymatic activity and protein abundance assays.

We next evaluated, for each map, how deleterious substitutions to each of the 20 possible amino acids tended to be. To this end, we calculated the median score for all substitutions observed for each of the 20 amino acids. For both maps, the correlation of this ‘deleteriousness vector’ with the corresponding vector derived previously from 28 other variant effect maps^43^ was strong (Figures S5A and S5B; Spearman’s R activity = 0.89, p<2e-16, R abundance = 0.89, p<2e-16), giving further evidence that the biochemical impact of changes to amino acids are captured. We also evaluated the impact of substitutions of the initial amino acids. The correlation of the aggregated ‘from’ amino acid scores to each the 28 other variant effect maps^43^ was moderate for the activity map and not significant for the abundance map (Figures S5C and S5D; Spearman’s R activity = 0.48, p=0.045, abundance p>0.05). We note that there is reduced power in the ‘from’ amino acid analysis given that many amino acids do not appear often in the reference sequence.

If the impact of missense variation on total enzyme activity was predominantly due to changes in protein abundance (e.g. arising through reduction of stability), we would expect high correlation between our total enzymatic activity and abundance maps. To assess this, we initially aggregated scores to and from each of the possible amino acids. We observed high correlation between abundance and activity maps when changing to possible amino acids, and moderate correlation when changing from the initial amino acid (Figure S5E and S5F; Spearman’s R to amino acid = 0.85, p<2e-16, Spearman’s R from initial amino acid = 0.41, p=0.01). Although, as might be expected given experimental measurement error, measuring correlation between abundance and activity maps using individual substitution scores yielded weaker correlation, it remained statistically significant (Figure S5G; Spearman’s R = 0.28, p<2e-16). The incomplete correlation we observed also supports the idea that deleterious SOD1 variants need not impact total activity via reduced protein levels, and many SOD1 missense variants likely affect specific activity but not abundance. However, for those variants exhibiting low abundance scores, we see a corresponding increase in the proportion of low activity scores (22%) relative to variants with high abundance scores (10%) (Figure S5H).

### 2.3 Initial evaluation of map quality

As an initial evaluation of the quality of our maps, we compared with the small-scale assays that had been used to validate the assays. For example, the likely pathogenic variant p.Ala146Thr, which our small-scale abundance assay found to be damaging (it was not tested in our small scale activity assay), appeared damaging in our abundance map. In another example, the loss of specific activity and abundance observed in small scale assays of the pathogenic variant p.Arg116Gly was experimentally confirmed in our maps. Finally, the “proxy benign” (PB) control p.Gly130Ser present in gnomAD that appeared tolerated in small-scale assays was also tolerated in both maps. There were no clear examples of discordance between large- and small-scale assay results except for p.Asn132Lys, which was found damaging in the small-scale activity map and not in the large-scale activity map.

As a first evaluation of the relevance of our maps to human biology, we investigated whether variants that appeared deleterious in our maps are selected against in the population. A previously-described missense constraint Z score^44^ derived from analysis of SOD1 variants in gnomAD was 1.2, indicating moderate selective pressure against these variants. In keeping with this, missense variants classified as damaging in our activity and abundance maps were less common in human cohort databases (see material and methods; total activity log-odds ratio = −1.1 and abundance log-odds ratio = −1.8; Fisher’s exact test; Figure S6). Thus, variants found damaging in each of our maps appear to be counter-selected in the human population.

Interestingly, a “loss-of-function intolerance” (pLI) score^44^ applied to gnomAD yielded a score 0.01 for SOD1, suggesting general tolerance to nonsense, and frameshift variants. This is consistent with the view that deleterious SOD1 variation is primarily due to its dominant gain of function impacts.

We next investigated, for both maps, trends related to amino acid substitutions with different properties. As expected, substitutions at hydrophobic residue positions had significantly more damaging effects in the activity map compared to polar and charged amino acids (Figure S7A; p_adj_=3e-9 and 1e-10, respectively, Wilcoxon test). Results for the abundance map were similar (Figure S7B; p_adj_=7e-30, 2e-37, respectively, Wilcoxon test). As expected, conservative substitutions at uncharged positions tended to be tolerated (higher scoring) in both maps relative to substitutions of other types (Figure S7C). Additionally, we found that proline substitutions tended to be damaging in both maps (Figure S7D; activity p_adj_=3e-8, abundance p_adj_=1e-7), as expected given the general tendency of proline residues to disrupt secondary structure.

### 2.4 Biochemical insights from SOD1 variant effect maps for activity and abundance

Substitutions that reduce both the net negative charge of SOD1 and also stability have been associated with lower survival time in ALS^45^. In keeping with this, the activity map showed that introduction of negatively charged amino acids tended to have a greater impact (lower score) than introduction of other amino acids (Figure S7E; Δmedian 0.08, p_adj_=7e-3), an effect that was even more strongly observed in the abundance map (Figure S7E; Δmedian 0.26, p_adj_=1e-3). This fits the previously proposed model that unstable SOD1 variants with maintained or modestly lowered net negative charge will tend to be more soluble, and therefore more likely to form toxic oligomers rather than accumulating as more benign larger aggregates^46^.

We assessed whether our scores agreed with known secondary structural features of SOD1 given the importance of core (buried) residues to protein stability (Figure S8; (ASA) < 20%). As expected, substitutions to core residues were poorly tolerated in both maps relative to non-homodimer surface residues (Figure 2A; Δmedian score between non-homodimer surface residues = 0.48, 0.99, p_adj_ = 2e-10, 2e-8, activity and abundance, respectively). Interestingly, the lowest-scoring category of residue positions was metal-binding positions for the activity map, and core positions for the abundance map. This supports the expected importance of metal binding for enzymatic activity of SOD1, while protein abundance depends strongly on the buried positions that would be expected to be important for maintaining stability.

**Figure 2.**
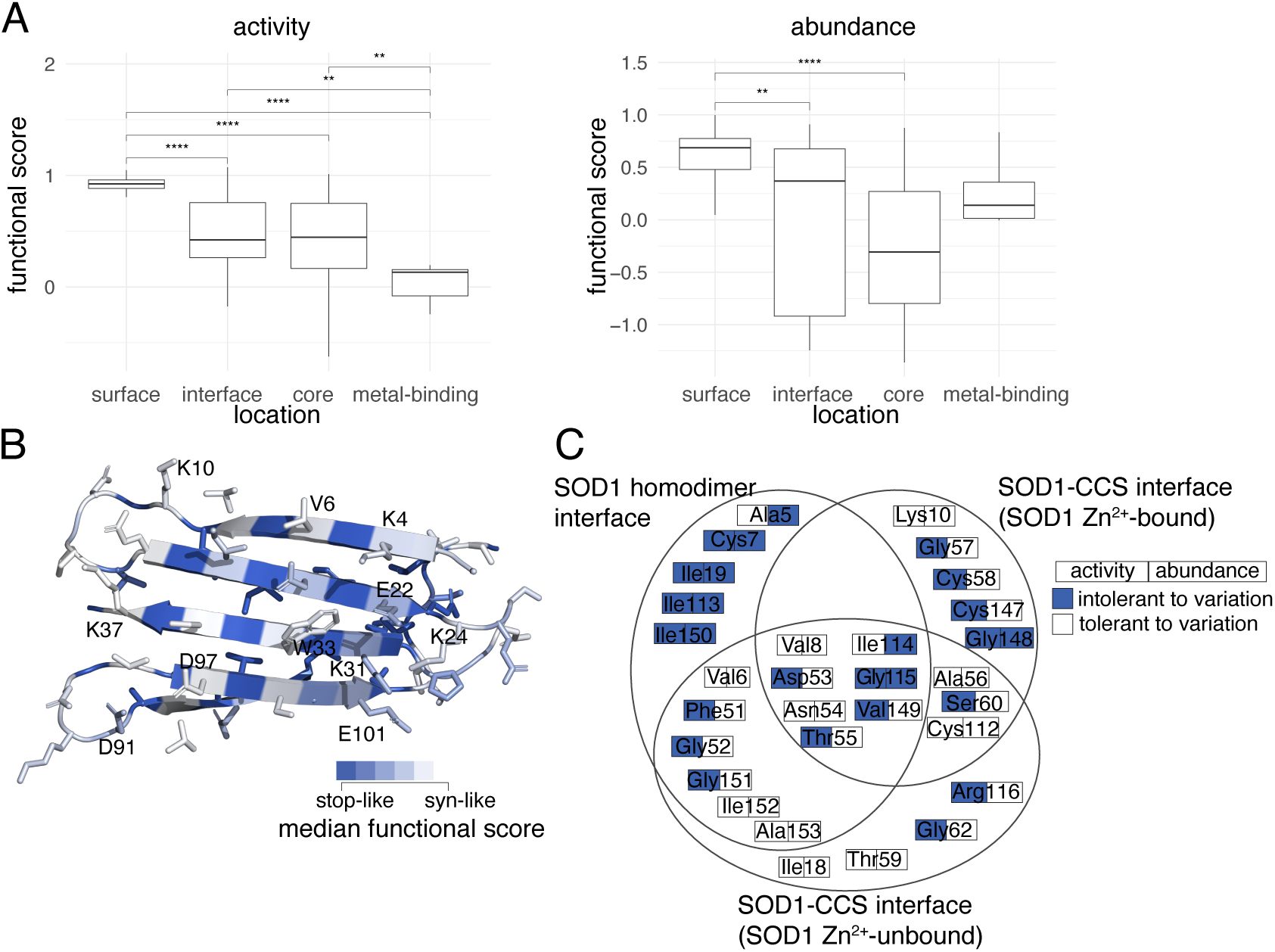
Modeling the effects of SOD1 missense variants on protein structure. (A) Distributions of median functional scores (where each median is of substitutions at each residue position) are shown for total enzymatic activity (left) and abundance (right) maps, summarizing sets of positions where the wild type residue is “core” (below 20% accessible surface area (ASA), “metal-binding”, “surface” (above 35% ASA), and “interface” (at the homodimerization interface). Boxes correspond to interquartile range, and bold bars indicate medians of the positional medians. Whiskers correspond to minima and maxima. P-values were calculated by Mann-Whitney U test. (B) Structural model of SOD1; colored according to the median functional score of substitutions at each position. (C) Venn diagram indicating whether amino acids found at SOD1 interfaces (either the homodimeric SOD1 interface or the SOD1-CCS interface with and without Zn²⁺-bound to SOD1) show for in each map whether the variant was tolerated (white) or intolerant (blue) to substitutions (see Methods for details on determination of threshold scores)

Hypothesizing that variants within the β-sheets of SOD1 would be enriched for stability impacts, we ‘painted’ a crystal structure of the active SOD1 homodimer^47^ according to the median score at each position for both maps (Figure S9A, S9B). Interestingly, residues intolerant to variation formed distinct ‘stripes’ on the β- sheets such that the intolerant positions tended to correspond to hydrophobic residues with inward-facing side chains in both maps.

To gain further insight into the mechanism of variant impacts, we examined activity and abundance map scores at specific residues of known biochemical importance in the context of the sequence of events known to promote SOD1 stability and activity. Briefly, this sequence begins with Zn²⁺ binding, Cu²⁺ binding, then associated release of CCS (“Copper Chaperone for SOD1”), formation of a disulphide bond between p.Cys58 and p.Cys147^48–51^, followed by formation of the active SOD1 homodimer.

We found that positions coordinating the binding of Zn^2+^ (p.His72, p.His81, p.Asp84), Cu^2+^ (p.His47, p.His49) or both metals (p.His64) were generally intolerant to mutations in both maps (Figure S10A), suggesting an impact on total enzymatic activity via an effect on protein abundance. Substitution of the disulphide-bonded p.Cys58 and p.Cys147 residues also showed severe functional defects in both maps. Interestingly, we found substitutions at the p.His121 residue (which has been implicated in Cu^2+^ binding^52^) impacted total activity but not abundance, suggesting a critical role in enzyme function that is independent of protein abundance. We expected that residues at the homodimer interface or within the core would be more sensitive to variation than non-interface surface residues (ASA > 35%). We expected that residues at the homodimer interface would be more sensitive to variation than non-interface surface residues (ASA > 35%). Indeed, for the activity and abundance maps, we found that substitutions at the homodimer interface exhibited more severe functional defects than other surface residues (Figure S2A; Δmedian=0.42; p_adj_=7e- 03, Δmedian=0.32; p_adj_=0.037 for activity and abundance map, respectively, by Mann-Whitney U test).

We next considered the electrostatic loop VII (residues 121–142), which contributes to a positively charged environment through which negatively-charged superoxide ions diffuse to the active site. Residues p.Lys137 and p.Thr138 within this loop, as well as the adjacent p.Arg144 residue, have been previously highlighted as important^53^. Our abundance map supported some contribution for p.Thr138 to stability but apparently not causing enough loss of total activity to affect fitness in our activity assay (Figure S10A). Residue p.Lys137 was tolerant to substitution in both maps. Interestingly, p.Arg144 was intolerant to substitution in the activity map, but not the abundance map, suggesting a stability-independent role in enzymatic function for this residue also.

Modifications, such as the oxidation of wild-type residues—specifically the solvent-exposed residues p.Val6, p.Trp33 and p.Cys112—have been linked to variant aggregation and to the propagation of gain of function effects^54–56^. Surprisingly, p.Val6, and p.Trp33 were both highly tolerant to substitution in both activity and abundance maps. Hydrogen bonding by p.Cys112 to both p.Glu50 and p.Arg116 residues has been noted as important to maintaining SOD1 structural integrity^57^. Also surprisingly, both activity and abundance maps indicated that, except for hydrophobic substitutions, p.Cys112 was quite tolerant to substitution (Figure S10B). Excepting hydrophobic substitutions, we found variation at p.Glu50 to also be generally tolerated, while both p.Arg116 and the p.Arg116-neighboring residue p.Gly115 (which may convey flexibility at p.Arg116) were intolerant to mutation.

It has previously been hypothesized for the aggregation-prone amyloid beta protein that specific key apolar residues can prevent aggregation by attenuating local hydrophobicity (the “amyloid stretch” hypothesis)^58^. Our analysis revealed that the charged p.Glu22 and p.Lys31 residues are intolerant to change in the abundance map (Figure S10C), which would be consistent with the amyloid stretch hypothesis if loss of attenuating effects of these residues on neighboring hydrophobic residues p.Trp33 and p.Val6 resulted in aggregation.

There are two previous ‘β-edge’ hypotheses related to the impact of charged residues at the edges of a β- sheet. First, it has been hypothesized (for β-sheet-containing proteins in general) that lysines located on edges of otherwise-aggregation-prone β-sheets are important for protein solubility^59^. Second, negatively charged residues on β-sheet edges have been described as gatekeepers preventing aggregation^60^. In contrast to the lysine β-edge hypothesis, and despite the fact many of SOD1’s lysine residues are located on the edges of the N-terminal β-sheet (more specifically p.Lys4, p.Lys10, p.Lys24, p.Lys31, and p.Lys37), all of these lysines appeared tolerated in our activity map, and all but p.Lys31 were tolerated in the abundance map (Figure 2B, Figure S10C). Our data is more consistent with the negatively-charged β-edge hypothesis, in that most of the negatively-charged residues on the β-sheet edge (p.Glu22, p.Asp91, and p.Glu101, but not p.Asp97) appear intolerant to variation in our abundance map. Overall however, the observation that all charged positions at the edge of a β-sheet appeared highly tolerant to variation in the total enzymatic activity map keeps us from drawing black and white conclusions about the consistency of our results with either of the β-edge hypotheses.

In addition to analyzing our scores in the context of the active SOD1 homodimer structure, we also considered SOD1 complexed with its paralog, the Copper Chaperone for SOD1 (CCS)^61^. We examined both a reported Zn^2+^-bound SOD1-CCS structure^61^ and a Zn^2+^-unbound SOD1-CCS model, derived via rigid-body docking of individual SOD1^62^ and CCS^63^ crystal structures (see Materials and Methods). In both SOD1-CCS docked models, we found residues in SOD1’s GDNT motif (positions 52-55) to be involved in CCS binding as previously reported^61^. We additionally identified a ‘Zn^2+^-SOD1-CCS’ set of five residues (see Figure 2C) as being involved in binding for the Zn^2+^-bound SOD1-CCS structure but not the Zn^2+^- unbound SOD1-CCS structure, suggesting their transient involvement in CCS-binding. Both GDNT and the Zn^2+^-SOD1-CCS residue set were found to be generally intolerant to substitutions in our activity map (Figure 2C). Four of the five Zn^2+^-SOD1-CCS residues also exhibited tolerance for substitution in our abundance map. Because CCS shares over 50% sequence identity with SOD1^64^, we might expect a CCS-SOD1 binding mode that closely resembles that of the SOD1 homodimer. However, only thirteen of the eighteen residues at the SOD1 homodimer interface are seen at the SOD1-CCS binding interface. Among these thirteen ‘both-interface’ residues, the activity map identified seven as intolerant to substitution—including three of the GDNT residues (p.Gly52, p.Asp53 and p.Thr55) and the GDNT-adjacent p.Phe51 residue— while the abundance map implicated only p.Ile114 and p.Gly115 as being important. One possible explanation for the discordance between the maps is the strict requirement for CCS1-dependent SOD1 metal-binding in yeast, while a CCS-independent pathway is sufficient for SOD1 metal-binding in mammals^65,66^.

### 2.5 Exploring mechanisms of active-site-adjacent substitutions

One way to gain insight about variant mechanism is to generate predictions of protein thermodynamic stability (ΔΔ*G*) for each variant. Agreement with functional scores immediately suggests a loss-of-stability mechanism. By contrast, ‘stable-but-inactive’ variants–defined by having a functional impact that cannot be explained by an impact on stability–are likely to act by other mechanisms. Across a range of proteins, such variants have been found to be substrate binding sites, surrounding ‘second-shell’ residues that influence these sites^41,67,68^, or to be important for protein dynamics^41,67,68^. Not surprisingly, ΔΔ*G* scores showed significant (albeit modest) correlation with functional scores (Figure S11A and S11B; Spearman’s R for activity = 0.35, p_adj_ = 5e-05; for abundance, R = 0.39, p_adj_ = 5e-06). To identify protein regions enriched for stable-but-inactive variants, we performed moving window analyses comparing activity and abundance map scores with predicted ΔΔ*G* values across all protein positions (Figure S11C). Although this revealed no broad regions enriched for stable-but-inactive variants, we next performed this comparison at each individual residue position.

Comparing the median scores for each map at each position with the median ΔΔ*G* value at that position identified several second-shell residues adjacent to the active site—p.Gly86, p.Asn87, and p.Gly128—as important for enzymatic function but not stability (Figure 3A). We first examined p.Gly86 and p.Asn87 within reported SOD1 structures, and hypothesized that one of these residues, p.Asn87, forms a salt bridge with p.Asp125, thereby stabilizing the local electrostatic field and facilitating a hydrogen bond network between p.Asp125 and the metal-binding residues p.His47 and p.Asp72.

**Figure 3.**
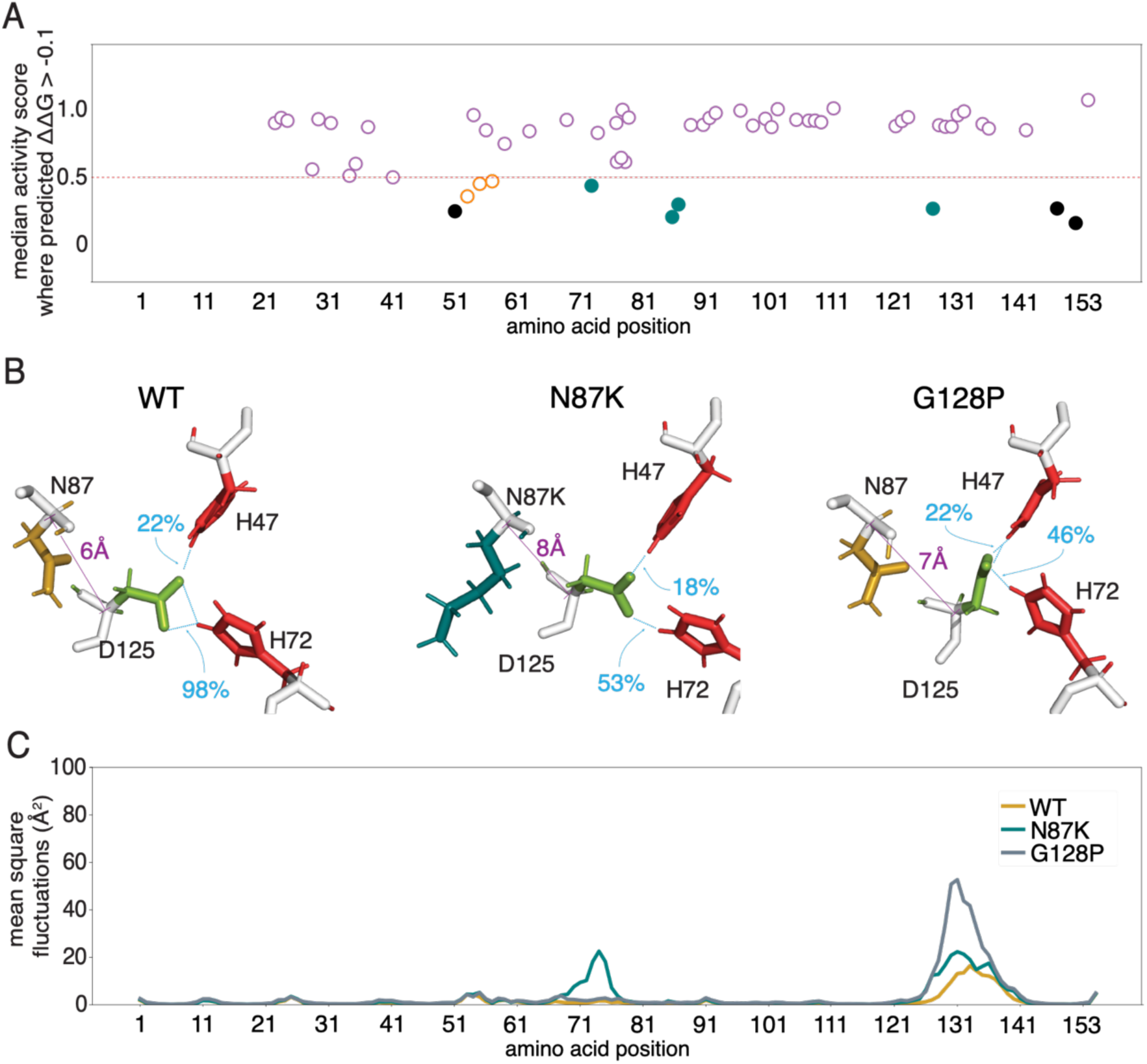
Modeling effects of SOD1 missense variants on protein stability and structure. (A) For the subset of SOD1 missense variants with no predicted detrimental impact on stability (median predicted *ΔΔG* < −0.1), median total enzymatic activity score is plotted. Filled circles indicate residues: a) at the SOD1-CCS interface (orange); b) residues proximal to the active site (‘second-shell residues’; teal); c) at the SOD1 homodimeric interface (black). (B) To illustrate the structural impact of variants Asn87Lys and Gly128Pro on the metal-binding site, the average distance (Å) between the Cα atoms of residue pairs 125 (electrostatic loop) and 87 (βstrand residue proximal to the electrostatic loop) is shown in purple, along with the fraction of simulation time (blue) in which hydrogen bonds occurred between metal-binding residue pairs Asp125-His47 and Asp125-His72. (C) Mean-square fluctuation (MSF) of Cɑ atoms of WT SOD1, as well as Asn87Lys and Gly128Pro variants. MSF reflects the average deviation of atoms throughout the simulation (400 ns) relative to the initial structure.

Based on MD simulations, previous studies concluded that folding of the electrostatic loop excludes water and promotes stabilizing hydrogen bonds between p.Asp125, p.His47 and p.His72^69^, together allowing SOD1 to bind both Cu^2+^ and Zn^2+^ ions. The electrostatic loop position can be understood in terms of hydrogen bonding of p.Asp125-p.His47 (indicating amenability to Cu^2+^ binding) and of p.Asp125-p.His72 (indicating amenability to Zn^2+^ binding)^69^. To quantify the relationship between the p.Asn87-Asp125 interaction and electrostatic loop position, we evaluated Cα distances for p.Asn87-Asp125 and both of the residue pairs that indicate amenability to metal-binding, using 400 ns MD simulations for both WT SOD1 and two SOD1 variants:

For WT SOD1, the Cα distance for p.Asn87-Asp125 was 6.4 ± 0.2 Å (Figure 3B), consistent with salt-bridging. Distances for p.Asp125-p.His47 and p.Asp125-p.His72 were 9.3 ± 0.3 Å and 10.2 ± 0.2 Å respectively (Figures S12A, S12B), providing a baseline WT SOD1 position for the electrostatic loop. Our observations were consistent with previous observations that WT SOD1 shows occasional hydrogen bonding for p.Asp125–p.His47 (22% of simulation time, reflecting occasional Cu²⁺-binding ability; Figure 3B, S12D)^69^. We observed constitutive hydrogen bonding for p.Asp125–p.His72 (98% of simulation time), reflecting stable Zn^2+^-binding ability (Figure 3B, S12E).

For SOD1 protein with the pathogenic p.Asn87Lys variant, we saw an increased p.Asn87-p.Asp125 distance of 7.6 ± 1.0 Å (Figure 3B), suggesting weakened electrostatic interaction. Mean distances for p.Asp125-p.His47 and p.Asp125-p.His72 increased to 9.9 ± 0.6 Å and 11.2 ± 1.4 Å (Figure S12A, S12B), with the electrostatic loop remaining in the Cu^2+^-binding state for 18% and in the Zn^2+^-binding state for 53% of the simulation time (Figure S12D, S12E). Further analysis of Cα atoms across all residues showed higher mean square fluctuations (MSF) values for this SOD1 variant, especially for residues near p.His72 and within the electrostatic loop (Figure 3C). More importantly, these results suggest that p.Asn87Lys impacts Zn^2+^-binding but not Cu^2+^-binding.

Although we did not specifically investigate the role of the p.Gly86 residue, given that this position is adjacent to p.Asn87 and that glycine is known to provide flexibility, we speculate that p.Gly86 provides the necessary conformational flexibility for p.Asn87’s role.

Returning to the second-shell residue p.Gly128 which showed intolerance to variation in the activity map that could not be explained by predicted stability effects, we hypothesized that this residue provides the flexibility needed for p.Asp125 to bond with the metal-binding sites. To further explore this, we simulated the dynamics of p.Gly128Pro variant SOD1. MD simulations showed that the p.Gly128Pro variant exhibited Cα distances of 7.1±0.5 Å, 9.7±0.5 Å, and 10.4±0.6 Å for p.Asn87-p.Asp125, p.Asp125-p.His47, and p.Asp125-p.His72, respectively (Figures 3B, S12A and S12B), with the structure remaining in the Cu²⁺- binding state for 22% (as seen for WT SOD1), and Zn^2+^-binding state for 46% of the simulation time (Figures S12D, S12E). Interestingly, unlike the p.Asn87Lys variant, the p.Gly128Pro structure displayed increased MSF values only within the electrostatic loop. More importantly, as observed for p.Asn87Lys, these results suggest that p.Gly128 impacts Zn^2+^-binding but not Cu^2+^-binding.

Taken together, these MD simulation results (summarized in Supplemental Table 1) suggest that p.Asn87 binding to p.Asp125 form a salt bridge that plays an important role in the positioning of the electrostatic loop and thus binding of both Cu^2+^ and Zn^2+^ ions. Our studies also support a previous suggestion that p.Gly128 introduces steric restrictions in the backbone of the electrostatic loop that create a barrier to the formation of metal-binding sites^69^.

### 2.6 Functional scores suggest genotype-phenotype association in SOD1-ALS

The impact of a given variant can manifest differently in different individuals—i.e., can have variable expressivity or incomplete penetrance—due for example to environmental influences that can include pathogen exposure, age, sex, and other factors. This variability impacts disease phenotypes, such as earlier disease onset or duration^37,70,71^.

We further investigated two pathogenic variants p.Asp12Tyr and p.His47Arg which are considered ‘clinically mild’ in the sense that onset of ALS is late and progression is slow^72–74^. While p.Asp12Tyr was non-damaging in both maps, p.His47Arg was non-damaging in the abundance map and found damaging in the activity map. The p.His47 residue, which binds Cu^2+^ directly, is both indirectly important for Zn^2+^ binding and required for SOD1 activity^75^. However, Cu^2+^ can still bind p.His47Arg variant SOD1 (at an alternative p.Cys112 site)^76^, potentially explaining why p.His47Arg is stable in our abundance assay, despite loss of enzymatic activity.

We next evaluated correlation more generally between our map scores and a previously-assembled collection^37^ of molecular and ALS disease phenotypes, including enzymatic activity and abundance in patient samples, protein half-life (of the human protein measured in mouse), and age of onset and duration of ALS disease. We saw significant correlation with SOD1 half-life for our abundance scores (Figure 4A; R=0.78, p_adj_=2e-2), but not for our activity map scores (p_adj_=1). We did not find significant correlation between either map and previous measurements of patient enzyme specific activity (Figure 4B; activity map p_adj_=1e-1; abundance map p_adj_=3e-1), nor with patient SOD1 protein abundance measurements (Figure 4C; activity map p_adj_=1e-1; abundance map p_adj_=1). However, scores for our abundance (but not activity map) showed significant correlation with age of onset (Figure 4D; activity map Spearman’s R=-0.13, p_adj_=3e-1; abundance map R=0.21, p_adj_=2e-2) while scores from both maps correlated significantly with disease duration (Figure 4E; activity map R=0.27, p_adj_=9e-3; abundance map R=0.34, p_adj_=5e-4).

**Figure 4.**
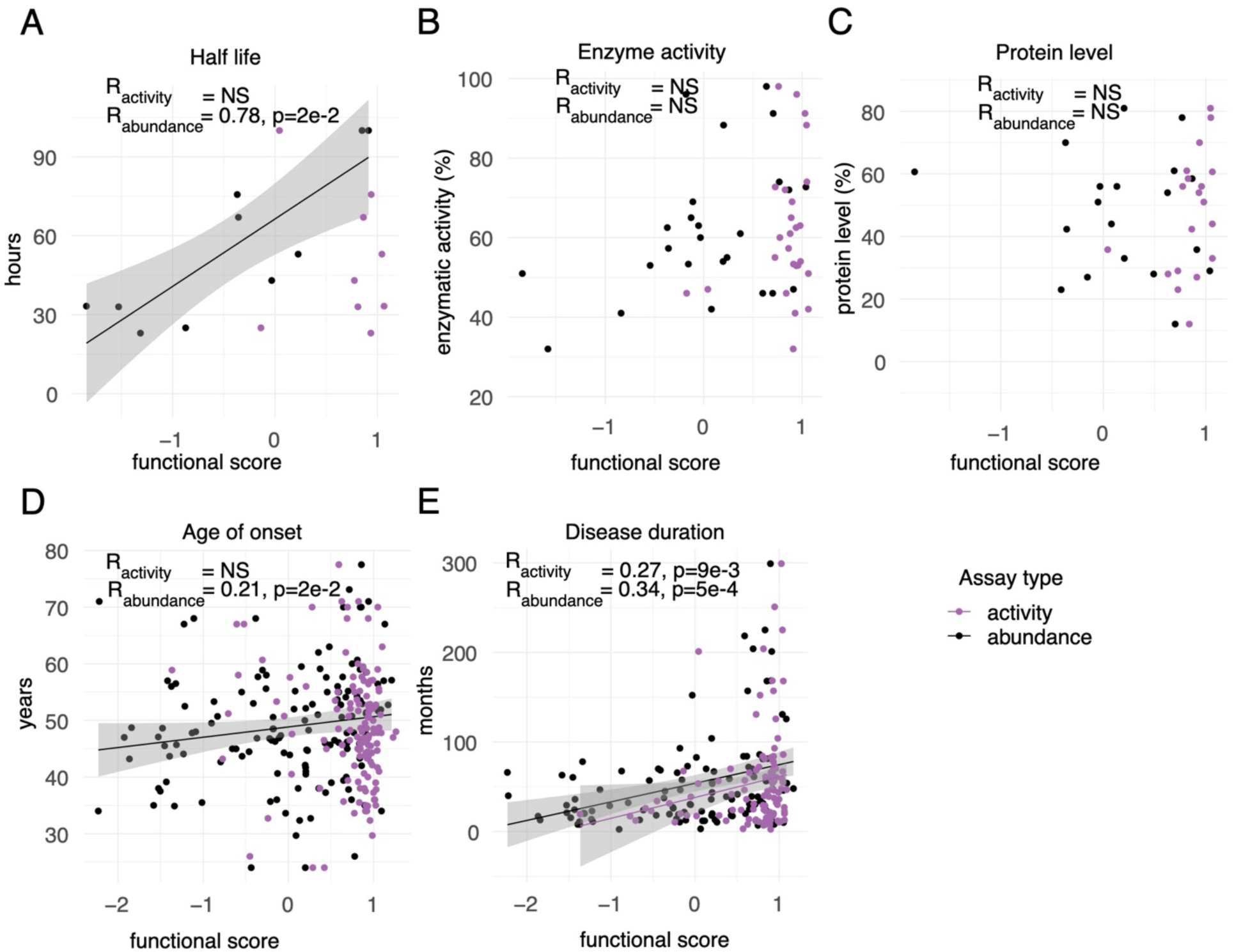
Genotype-phenotype analyses of SOD1 variants in both assays against a dataset of various phenotypes. Scores for both activity and abundance maps (mean score for substitutions at each position) were compared with an assembled dataset of patient phenotypes including (A) Mouse protein half-life measurements, (B) Patient enzyme specific activity, (C) Patient protein abundance measurements, (D) Age of ALS onset, (E) Disease duration^37^. NS = not significant. Grey indicates 95% confidence interval for significant correlations. See methods for details on regression and significance calculations.

Taken together, while these analyses revealed only limited correlation between our maps and previous biochemical data, they showed correlation with disease phenotypes, supporting both the quality of the maps and the existence of quantitative genotype-phenotype associations for classical ALS.

### 2.7 SOD1 abundance scores correlate with variant pathogenicity

Next, we wished to assess the ability of the SOD1 maps to identify pathogenic variants. More specifically, at a series of score thresholds for each map, we wished to evaluate precision (fraction of variants below a given threshold score that have been annotated as pathogenic or likely pathogenic (P/LP)) and recall (fraction of all variants annotated as P/LP that scored below the threshold). Because precision is impacted by the proportion of pathogenic variants in the reference set (which will not generally correspond to the proportions observed clinically), we adopted the ‘balanced precision’ measure which estimates the precision that would have been observed with a balanced reference set (see materials and methods).

To enable this analysis, we established two positive reference sets of P/LP variants: 85 observed by Labcorp Genetics in clinical sequencing^77^, and 42 high-confidence variants from ClinVar^12^. Obtaining a negative reference set was more problematic due to the lack of SOD1 missense variants annotated as “benign” or “likely-benign”^12,78^. We therefore selected 100 PB variants from gnomAD which have no annotated disease association or evidence of functional impact. We note that, to the extent that our negative reference set is contaminated with truly pathogenic variants, or our positive reference set contaminated with truly benign variants, our performance estimates will tend to be conservatively low.

Using the Labcorp Genetics and ClinVar positive reference sets, pairing each in turn with the PB negative reference set, we derived balanced precision vs recall curves for both our activity and abundance maps (Figure 5, Figure S13). From each curve, we derived two summary measures of performance: area under the balanced precision vs recall curve (AUBPRC) and recall at 90% balanced precision (R90BP). For the activity map, this analysis showed AUBPRC values of 0.58 and 0.56 for ClinVar and Labcorp Genetics analyses, respectively. By contrast, the abundance map showed relatively higher AUBPRC values of 0.79 and 0.80.

**Figure 5.**
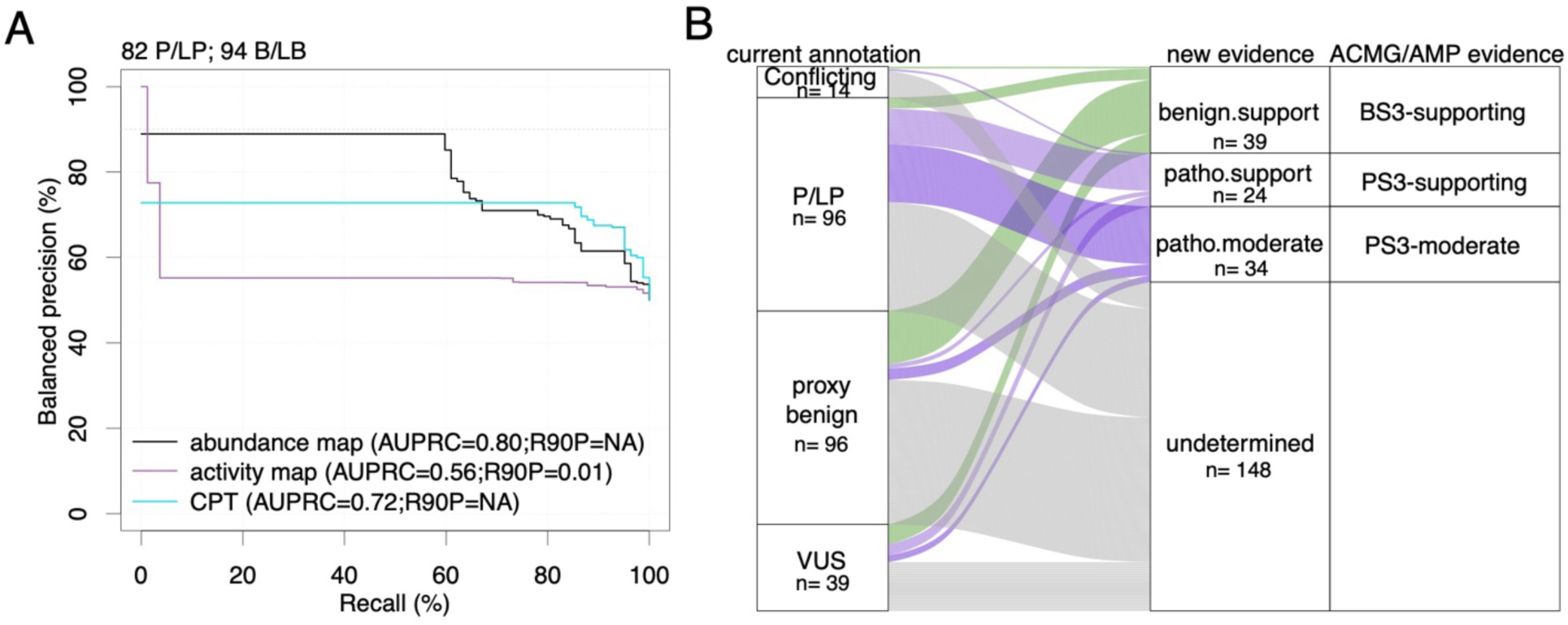
The SOD1 abundance map best distinguishes positive from negative reference variants, and provides evidence for 41% of VUS. Evaluation of precision (fraction of variants in the positive reference set which scored below each threshold functional score) vs. recall (fraction of positive reference variants with functional scores below threshold) shown in (A). More specifically, we used “balanced precision” such that precision values reflect performance in a setting where positive and negative sets contain the same number of variants for reference sets with positive variants obtained from Labcorp Genetics. Balanced precision vs. recall curves are shown for the SOD1 total enzymatic activity (purple) and abundance maps (black), as well as for the best-performing computational predictor CPT (turquoise). (Performance for three additional predictors is shown in Figure S13.) Performance was summarized in terms of area under the balanced precision vs recall curve (AUBPRC) and recall at a balanced precision of 90% (R90BP). Plots indicate the number of variants in the positive (P/LP) and gnomAD negative (PB) reference sets used here (see Methods for details). Currently clinically annotated variants and their annotations are shown in (B). New evidence proposed based on calibrating our abundance map scores against the ClinVar or Labcorp Genetics reference sets is shown in the right column. (See main text and Methods for details on the derivation of evidence strength).

It can be instructive to compare the performance of variant effect maps with computational predictors. Overall, we found that our abundance map outperformed four widely used predictors when assessed with ClinVar or Labcorp Genetics reference sets, respectively—AlphaMissense^79^ (Figure S13; AUBPRC = 0.70; 0.72), ESM-1b^80^ (AUBPRC = 0.72; 0.69), CPT^81^ (AUBPRC = 0.73; 0.72), and VARITY^82^ (AUBPRC = 0.68; 0.71). We were unable to calculate R90BP values for computational predictors, as none exceeded 90% balanced precision for either reference set at any recall threshold. Moreover, a moving window analysis did not reveal any SOD1 subregion for which computational predictors showed substantially better performance (Figures S14).

To provide our map in a form that is convenient for use in interpreting SOD1 missense variants, we used an established kernel density estimation method^83,84^ to transform all abundance map scores into log-likelihood ratios of pathogenicity (LLRp; see Methods) for the Labcorp Genetics positive reference set (given that there were more reference set variants available than in ClinVar) and PB negative reference set (Figure S15). To calibrate these scores for use within the ACMG/AMP framework for variant interpretation, categorical evidence strength levels were then derived from LLRp values as previously described^41,83^. Although these estimates for evidence strength were obtained using an objective empirical framework, we caution that this procedure has not yet been reviewed by the appropriate ClinGen variant curation expert panel or equivalent clinical body. That said, of the 41 SOD1 missense variants currently annotated as VUS (Figure 5B), the abundance map provided scores for 39 of them and yielded supporting evidence toward benign for 9 (22%), supporting evidence towards pathogenic for 5 (12%), and moderate evidence toward pathogenic for 3 (7%). Thus, our map provides new evidence for over 41% of existing VUS, offering the potential to improve sensitivity and accuracy of SOD1 variant interpretations.

## Discussion

Using a combination of functional assays that respectively reflect total enzymatic activity and protein abundance, we generated proactive variant effect maps evaluating the functional impact of nearly all missense variants of human SOD1.

Functional scores for both maps largely recapitulated known sequence-structure-function relationships for SOD1. For example, buried positions scored significantly lower than non-interfacial surface positions on both maps. Additionally, substitutions at hydrophobic positions were significantly more damaging to both activity and abundance, while conservative substitutions at uncharged sites were better tolerated. Moreover, proline substitutions were generally damaging in both maps, aligning with proline’s known role in disrupting secondary structure. In addition to these patterns of mutational tolerance, we found missense variations to be damaging at positions crucial for metal ion (Zn^2+^ and Cu^2+^) binding and the formation of the p.Cys58-p.Cys147 disulfide bond. Variation at positions of residues required for Cu-dependent CCS- mediated activation of SOD1 (i.e., Cu^2+^ insertion and intra-subunit disulfide bond formation) were well tolerated in our human cell-based abundance map, aligning with previous reports of a CCS-independent pathway of SOD1 activation in human cells^85^. These positions were relatively intolerant to variation in the yeast-based activity map, consistent with the importance of CCS-dependent SOD1 maturation in yeast^66^.

Our maps provide supporting evidence for three existing hypotheses. First, the maps showed that introduction of negatively charged amino acids tended to be detrimental, consistent with the hypothesis that such substitutions increase the toxic, soluble form of SOD1 and correlate with reduced survival in SOD1- ALS patients^45^. Second, our finding that substitution of solvent-facing hydrophobic residues with charged residues was not tolerated supports the ‘amyloid stretch’ hypothesis, which posits that charged residues mitigate the aggregation-prone effects of adjacent hydrophobic residues^58^. Third, our maps showed that substitutions in charged residues on the edges of β-sheets (p.Glu22, p.Asp91 and p.Glu101), tended to be damaging, supporting a previous hypothesis that positioning of charged residues at β-sheet edges decreases aggregation propensity^59^. Although primarily cytosolic, SOD1 is also secreted. SOD1 variants reportedly require a certain degree of solubility for aggregation and prion-like propogation ^86^. It would be interesting and clinically valuable to investigate impacts of p.Glu22, p.Asp91 and p.Glu101 on initiating aggregation or prion-like propagation of misfolded SOD1.

Our map scores suggested a new hypothesis, that the interaction between p.Asn87 and p.Asp124 (as part of the electrostatic loop) stabilizes a ‘closed’ conformation that excludes solvent molecules from the metal site and enhances Zn²⁺ and Cu²⁺ affinity by maintaining strong hydrogen bonds between p.Asp124 and residues p.His46 and p.His72. Consistent with previous reports^69^, our MD simulations also supported a hypothesis that substitutions at p.Gly128 alter the flexibility of the electrostatic loop, disrupting its proper positioning, increasing solvation, reducing interaction with the Zn²⁺-binding p.His72 residue, and impairing overall protein stability. Further studies to confirm the validity of our MD simulation results might focus on investigating the role of the p.Asn87-p.Asp125 interaction in controlling the flexibility and positioning of the electrostatic loop, e.g., our map and MD simulations predict that *in vitro* enzymatic and agitation-induced fibrillation studies of the p.Asn87Lys variant would show impacts on both protein activity and stability.

Our abundance map scores were significantly correlated with disease duration and age of onset in SOD1- ALS patients, supporting a genotype-phenotype relationship. Unexpectedly, abundance map scores did not correlate with patient SOD1 protein abundance measurements, despite showing strong positive correlation with mouse protein half-life^37,87^; Low SOD1 protein half-life has been attributed to increased chaperone-assisted autophagy, leading to chaperone exhaustion, higher levels of misfolded SOD1, and ALS onset^88,89^. A possible explanation for the observed *negative* correlation of the abundance map with patient/mouse abundance measurements is homeostatic upregulation of SOD1 expression in SOD1-ALS patients, particularly in nervous system tissue^90,91^. Although such homeostasis could compensate for reduced protein half-life *in vivo*, we should not expect to observe this in the exogenous (TetOn) expression system we used for our assays. Support for genotype-phenotype concordance was also demonstrated by variants p.Asp12Tyr and p.His47Arg—deemed pathogenic by ACMG/AMP evidence criteria but associated with a mild clinical course of ALS characterized by late onset and slow progression^22^ were found to be tolerated in our maps. With the increasing routine use of clinical genetic testing, future studies could explore whether abundance impact scores are predictive of disease severity and, more specifically, whether high abundance correlates with clinical benignity.

Our maps revealed that residues at the homodimerization interface are highly sensitive to variation, supporting the idea that dimerization is required for SOD1 activity^92^. That our abundance map reflects the importance of abundance to disease is supported by: the ∼0.1 nM *Kd* for SOD1 dimer dissociation^93^, the significant correlation observed between our abundance map scores and reported protein half-life measurements, and the ability of the abundance map to identify pathogenic variants. Indeed, maintaining lower concentrations of the SOD1 monomer has been shown to prevent its prion-like propagation^94^.

Our variant effect map scores correlated with pathogenicity, as measured via precision vs recall analysis. The best performance came from the abundance map, surpassing that of top-performing computational predictors such as AlphaMissense, CPT, ESM-1b, and VARITY.

An important limitation of our analysis is that no clinical missense variants of SOD1 are currently classified as benign, so that our negative reference set was necessarily composed of ‘‘proxy benign’’ variants from gnomAD. Moreover, such contamination should similarly impact performance of both our map and computational predictors, except perhaps to the extent that computational predictors have been overfit to the pathogenicity labels of variants used in our reference sets.

Another caveat of our study is that measurements were subject to random error. For example, a minority of nonsense variants identified as damaging in yeast-expressed cDNA were not detected as damaging in the human-cell-based VAMP-Seq assay. Although we used previously described methods^39^ to evaluate random errors associated with each experimental score, reflecting the estimated magnitude of random experimental error, this does not prevent random errors that move in the same direction for two replicates for some small subset of variants.

Our measurements may also have been subject to systematic errors from various sources. For instance, our variant pools contain clones with multiple variants, which TileSeq cannot detect if they are outside the sequenced tiles. Such unseen variants can include mutations in the fused GFP construct. Indeed, in the human pre-selection library, approximately 10% of the variants showed a loss of GFP, with a recurrent deletion breakpoint in the linker region between SOD1 and GFP. That each observed variant is present in many independent clones will tend to mitigate the effects of secondary, unseen variants within any given clone, and the separation of scores for synonymous and nonsense variants in both libraries suggest that influences on any given variant’s score by unseen distal variants do not represent a major issue.

Another caveat is that our activity map evaluated protein function within the context of a yeast cell. As noted above, the yeast assay may have erred in finding residues as intolerant to substitution because of their role in facilitating CCS-dependent SOD1 maturation in yeast, given the CCS-independent nature of SOD1 maturation in human cells. In addition, the yeast-based activity map failed to detect the functional impact of many variants outside the active site, including some variants previously shown to induce unstable (yet active) SOD1 that also yield toxic gain of function (aggregation) typical of SOD1-ALS. We speculate that the yeast chaperone Hsp104, which lacks a mammalian or vertebrate homolog, may have restored solubility of some mutant SOD1 protein, thereby increasing activity and cell growth in our assay^95,96^.

We also acknowledge that both activity and our (human cell-based) abundance assays may have missed other potential SOD1-related ALS pathomechanisms. For example, gain- and loss of function effects of SOD1 variants have been linked to the prion-like propagation of the misfolded protein along neuroanatomical pathways^97,98^. Although our abundance map provides evidence to identify pathogenic variants, future studies might assess the cell-non-autonomous ability of SOD1 variants to initiate or sustain prion-like propagation of misfolded conformations across a broad neuronal lineage.

A further limitation of our maps is their inability to detect aggregation (except indirectly to the extent that it impairs activity or abundance). SOD1 variants associated with toxic gain of function aggregation in cell lines have been linked to the accumulation of non-native oligomers, particularly the metastable SOD1 trimer^99^. Aggregates of SOD1 have also been widely reported in ALS patient samples and animal- and cell-models of the disease^100^. However, the reported aggregating phenotype of SOD1 variants in cell-based models can be attributed to overexpression, leading to spontaneous non-native folding intermediates, stoichiometric exhaustion of the chaperone pool, or both. This is supported by our failure to observe robust aggregation except in the context of both transient overexpression of WT or p.Ala5Val, p.Gly86Arg variants of SOD1 and treatment with a protease or heat shock protein 70 inhibitor. For cells stably expressing variants, known gain of function variants showed reduced scores in our abundance map, suggesting the degradation of misfolded SOD1^101^. Given the genetic heterogeneity of ALS, further studies should focus on the gain of function effects of SOD1 variants, especially in the context of impairment of molecular chaperones, which are not only important for re-folding but also for promoting the degradation of misfolded proteins^101^.

Other contexts to explore that might modulate SOD1 variant effects include redox state, which has been suggested to affect the propensity of SOD1 to aggregate^102,103^. In pilot studies, we found the aggregation propensity of SOD1 variants to be indistinguishable from WT. However, a human-cell-based complementation assay with the potential to capture more tissue-specific mechanisms might increase sensitivity to aggregation. It would also be interesting to investigate potentially disease-accelerating functional effects of variants in neuronal cell models, across different environmental contexts such as oxidative stress.

Our observation of a variant being deleterious—whether in the context of our enzymatic activity or protein abundance assays—does not, by itself, provide a full mechanistic explanation. For example, misfolding of SOD1 variants could contribute to toxicity through multiple, potentially overlapping pathogenic pathways. For example, misfolding could promote toxic oligomeric assemblies formed by gain of function pathogenic variants, but also contribute to degradation that limits the toxicity of these assemblies. Loss of function impacts (on either abundance, activity, or both) could reduce the enzyme’s protective role of limiting damage from free radicals in the nucleus ^99,100^. SOD1-related ALS pathomechanisms further include excitotoxicity, oxidative stress, mitochondrial dysfunction, and altered Ca²⁺ metabolism^100^. We note that our abundance assay relies on the fusion of SOD1 to GFP, which may slightly alter physicochemical properties of the protein^104^. Thus, while our abundance and activity map scores each demonstrate value towards identifying pathogenic variants, they should be used cautiously when inferring benignity. In the future, additional scalable assays could be employed to generate a more comprehensive pathomechanistic atlas of SOD1 variant effects.

The SOD1 variant effect maps we provide could have immediate value: For a symptomatic person with a SOD1 variant that would otherwise be classified as VUS, our functional evidence could help confirm an early ALS diagnosis that was otherwise lacking definitive support, thus enabling earlier therapeutic intervention and clinical trial enrollment. Identification of each additional pathogenic variant could also enable cascade screening to identify family members who are carriers, enabling genetic counseling and surveillance.

Finally, we note the recent appearance of a preprint describing a large-scale single-cell RNA-seq analysis of the effects of SOD1 missense variation^105^. Once a final peer-reviewed version of each study has appeared, it will be interesting to carry out a comparison and (perhaps) integration of these maps.

Beyond the immediate clinical benefits of offering robust evidence for stronger interpretation and reclassification of variants, these variant effect maps also offer a resource for deeper insights into sequence-function relationships for SOD1, contributing to the growing atlas of variant effects^106^.

## Materials and Methods

### SOD1 variant notation

Amino acid substitution notations (e.g. p.Ala5Val, p.Gly86Arg, p.Gly94Ala) correspond to the current SOD1 NCBI:NC_000021 code-determining nucleotide sequence.

### Cell lines, yeast strains and plasmids

The *Saccharomyces cerevisiae* strain (*MATa sod1Δ::KanMX his3Δ1 lys2Δ0 leu2Δ0 ura3Δ0*) used to assay activity for *SOD1* variant libraries was obtained from Horizon Discovery (formerly Open Biosystems). For yeast expression, WT and mutant SOD1 open reading frames (ORFs) were subcloned into the Gateway-compatible yeast expression vector pHYC-DEST2 (CEN/ARS-based, ADH1 promoter, *LEU2* marker)^107^. The SOD1 ORF clone (Ensembl: ENSG00000142168, GenBank: NM_000454.5) was obtained from the Human ORFeome v.9.1 library ^108^.

HEK293T T-REx Bxb1 landing pad cells, kindly provided by Dr. D. Fowler, were also used to assay SOD1 variant libraries. To generate the Gateway-compatible human integration vector pDEST-HC-REC-v2 for the Bxb1 landing pad, the attB-PTEN-IRES-mCherry plasmid (also a gift from Dr. D. Fowler) was obtained and the Kozak sequence and PTEN open reading frame replaced with the 1873 bp Gateway attR1-CcdB-attR2 cassette from pDEST-AD-CYH2. We used overlap extension PCR to generate a ‘SOD1 abundance reporter’ construct with GFP (Genbank: KC896843.1) fused to the C-terminus of SOD1.

Using Gateway LR reactions, both WT and variant disease-associated SOD1 ORFs were subcloned into the yeast expression vector (pHYC-DEST2), and both WT and variant versions of the SOD1-GFP construct were subcloned into the pDEST-HC-REC-v2 human integration vector. ORF identity and expected mutations were confirmed by Sanger sequencing. Yeast expression vectors (including an empty vector control bearing the ccdB marker under a bacterial promoter were transformed into the appropriate *sod1Δ* yeast strain. Human integration vectors (including an “mCherry only” negative control bearing an mCherry construct in place of SOD1-GFP) were transfected into HEK293T cells.

Although the phenomenon of wild-type (WT) SOD1 stabilizing SOD1 variants through heterodimerization has been reported^109^, this has not been implicated as a pathomechanism for SOD1-ALS^110^. However, to allow for this possibility, we did not knock out the endogenous *SOD1* locus in the HEK293T cell line we used.

### Validation of a yeast-based total enzymatic activity assay

From a single colony, *sod1Δ* yeast strain cells expressing human SOD1 wild-type (WT), empty vector control or variant plasmid were grown to saturation at 30°C. Each culture was then adjusted to an optical density at 600 nm (OD_600_) of 1.0 and serially diluted by factors of 5^1^, 5^2^, 5^3^, 5^4^, and 5^5^. These cultures (5 mL of each) were then spotted on SC-LEU plates as appropriate to maintain the plasmid and incubated at either 30°C or 38°C for 48 hours. After imaging, results were interpreted by comparing the growth difference between the yeast strains expressing human genes and the corresponding empty vector control.

### Small scale abundance assay validation in mammalian cells

Analysis of GFP expression profiles in transfected HEK293T T-REx Bxb1 landing pad cells was conducted for WT, empty vector control (mCherry+ only), and variants at day 7 post-transfection with plasmid and Bxb1 recombinase following 72-hour induction with 2 μg/mL doxycycline, using a Beckman-Coulter Gallios flow cytometer. Each population was gated for live (Sytox Red Dead Cell Stain) and singlet (FSC/SSC) cells, and subsequently for integrants based on being negative for BFP and positive for mCherry. Compensation was applied to correct for GFP fluorescence at the wavelength used for mCherry fluorescence detection.

### Generating mutagenic libraries

Libraries were generated using an updated version of Precision Oligo-Pool Based Code Alteration (POPCode) approach^40^ which eliminates a requirement of the previous POPCode approach^40^ for a uracilated template (See Document S1). Oligos were designed along the entire coding region of SOD1, such that each oligo contained an “NNK” degenerate code centered on each codon (see Table S1 for Primers). Although oligo designs differed only near the SOD1-GFP junction, for convenience we generated separate libraries for both SOD1 and SOD1-GFP. To generate the mutagenic libraries, the template plasmid backbone was denatured and the pooled, phosphorylated oligos were annealed along with primers that add an overhang ‘tag’ to allow for preferential amplification of the mutagenized strand by PCR. A fill-in reaction was performed with Phusion MM (NEB), and Taq DNA ligase (NEB) was applied to seal the nicks. Subsequent PCR reactions amplified the mutagenized strand using the tag sequences, and added attB1 and attB2 sites for Gateway cloning into destination vectors compatible for either yeast or human cell-based assays (see Table S1).

### Multiplexed assay for total enzymatic activity

Total enzymatic activity of individual SOD1 variants was examined using a yeast-based functional complementation assay: The mutagenic library without GFP was transformed into the *S. cerevisiae sod1Δ* strain using the EZ Kit Yeast Transformation kit (Zymo Research), and ∼1,000,000 transformants pooled to form the host library. Yeast transformants were grown at 30°C in synthetic complete (SC) media with glucose as the carbon source, lacking leucine (SC-LEU; USBiological) to ensure plasmid retention (pre-selection condition). Plasmid pools were prepared from 10 optical density units (ODU) of cells (defined as the number of yeast cells in 10 mL of a 1 OD600 culture, typically 10^8^ cells) and used as templates for downstream tiling PCR. Two replicates of approximately 4×10^8^ cells from the transformant pool were each inoculated into 200 mL SC-LEU medium and grown at restrictive temperature (post-selection condition; 38°C) for 48 h. In parallel, the *sod1* mutant strain was transformed with the WT ORF and grown alongside the mutagenic pool. Plasmids were extracted from 10 ODUs of cells from each culture (two replicate cultures for both post-selection condition and WT control) and used as templates for downstream tiling PCR (see Table S1).

### Large-scale mammalian assays

The impact of SOD1 variation on protein abundance was evaluated in a human cell line using the VAMP-seq method^35^. The mutagenic SOD1-GFP library and Bxb1 recombinase in a 10:1 mass ratio was transfected using FuGene 6 into two replicates of approximately 15×10^6^ HEK293T Bxb1 landing pad cells, enabling the expression of a single variant per cell. After induction with 2 ug/mL doxycycline, ∼2M cells per replicate were sorted 7 days post-transfection using a SONYMA900 cell sorter, gated for live cells (those unstained with SYTOX Red dead cell stain), singlets based on FSC/SSC, and enrichment for BFP- and mCherry+ cells. Integrated cells were expanded for 7 days and ∼2M cells sorted on the SONYMA900 for the post-selection condition: high-GFP and high-mCherry (high transcript and high expression). The pre-selection condition was 2M cells re-sorted for integration with the mCherry marker on the same day as the post-selection condition. Cells of the pre-selection and post-selection conditions were expanded for 7 days and pellets of 1×10^7^ cells were collected. gDNA from 2 pellets for each biological replicate condition was extracted with the Sigma gDNA extraction kit. All of purified DNA was amplified using primers BxbAmp_Tet_F2 and Bxb1Amp_R1 (see Table S1) and used as templates for downstream tiling PCR.

### Sequencing and large-scale analysis

For plasmid (total enzymatic activity) and amplicon (abundance) libraries from both pre-selection and post-selection conditions, five short template amplicons (∼150 bp) that tile the SOD1 ORF were amplified with primers carrying a binding site for Illumina sequencing adaptors (see Table S1). In a second-round PCR, Illumina sequencing adaptors with index tags were added to the first-step ‘tiling’ amplicons. Paired-end sequencing was conducted on all tiles, significantly reducing base-calling errors and enabling accurate detection of very low (parts-per-million) variant frequencies. Each assay was independently sequenced using an Illumina NextSeq 500 with a NextSeq 500/550 High Output Kit v.2, achieving a sequencing depth of >1,300,000 paired-end reads per tile. Sequencing reads were demultiplexed with bcl2fastq v.2.17 (Illumina).

Raw read data was processed and functional impact (total enzymatic activity or abundance) scores for variants were assigned as previously described ^40,41^. Briefly, we quantified variant allele frequencies using TileSeq_MutCount (https://github.com/RyogaLi/tileseq_mutcount), which incorporates Bowtie2 for aligning paired sequencing reads to the reference template. The posterior probability for each divergent base-call was determined, and those surpassing the 0.9 threshold were counted. We then filtered out variants with read counts less than 10 or frequencies below the 90th percentile of the WT. Next, we generated a score for each variant using the tileseqMave (https://github.com/jweile/tileseqMave) pipeline. After calculating the error-corrected enrichment ratios by subtracting WT variant frequencies from both the pre-selection and post-selection libraries, a functional impact score was determined for each variant based on its relative enrichment compared to the median ratios of nonsense and synonymous variants. Scores were then rescaled such that the medians of nonsense and synonymous variants were 0 and 1, respectively. Biological replicate scores were averaged and error calculated using an inverse-variance weighted average of scores, with weights proportional to the combined standard deviation computed from the weighted variances and degrees of freedom. Use of inverse-variance weighted averages provided greater weights for scores with lower uncertainty. For variants with no replicate, the standard error and ‘degrees of freedom’ were taken directly from the existing biological replicate.

### Reference sets and benchmarking

To initially evaluate the ability of total enzymatic activity and abundance assays to predict variant pathogenicity, we curated a small reference set of pathogenic/likely-pathogenic variants from ClinVar selecting those that spanned the length of the SOD1 coding sequence. “Proxy-benign” p.Asn20Ser and p.Gly130Ser variants were chosen based on their presence in gnomAD with no corresponding annotation.

To assess the ability of variant effect maps to identify pathogenic variants in large scale, we then used a “positive” set of 85 and 42 variants annotated as pathogenic/likely pathogenic (P/LP) based on Labcorp Genetics’s variant interpretation framework (Sherloc)^77^ and ClinVar P/LP variants with multiple submitters^12^, respectively. Due to the absence of reported benign variants, we augmented the negative set with 100 “proxy-benign” variants from gnomAD v.4.1.0, requiring only that these variants have no reported clinical annotations. To assess the performance of the maps and computational predictors of disease variants, we evaluated the trade-off between precision (proportion of variants below a threshold that are pathogenic) and recall (fraction of known pathogenic variants identified)^42^. To account for reference set size imbalances and varying prior probabilities, we generated “balanced” precision-recall curves which weigh the positive and negative reference sets 50/50 using a Bayes’ Rule-based method adjusting precision for prior probability^82^.

To determine whether variants were tolerant or intolerant to variation in either the activity or abundance map, we set a threshold of scores < 0.5 for the activity map and <-0.4 for the abundance map, with cutoffs set based on the minimum frequency of variants located between the bimodal distribution of missense variant scores.

### Transformation of scores to log-likelihood ratios of pathogenicity

To provide a quantitative Bayesian evidence weight for clinical variant interpretation, we estimated the log-likelihood ratio of pathogenicity (LLRp) for each variant’s abundance score. (This analysis was not performed for the yeast activity map given poor precision vs recall performance.) To this end, probability density functions were separately estimated for positive and negative reference variant score sets using kernel density estimation (more specifically, the Epanechnikov kernel^111^ with bandwidth determined by the Sheather and Jones method^112^). For variant scores in each map, a pathogenic:benign log-ratio was calculated using the estimated probability density functions. LLRp values were then calibrated against thresholds established by the ACMG/AMP variant classification system, using an adapted version ^41^ of the calibration approach developed by Tavtigian *et al*^83^.

### Thermostability calculations

Towards determining whether variant impacts in our total enzymatic activity map could be attributed to changes in steady-state protein level (abundance), as opposed to specific activity, we calculated protein thermostability (*ΔΔG*) values. Calculations of *ΔΔG* were obtained from the stability predictor DDGun3D version 0.0.2^113^ The PDB entry 1HL5^47^ for SOD1 was to predict these values, and satisfied the following conditions: an X-ray determined structure with resolution 1.8 Å or better, and no missing or non-standard residues. We defined stabilizing amino acid substitutions as those for which *ΔΔG* >0, and a destabilizing substitution for DDG < −0.5. Scores obtained from DDGun3D are available in Table S1).

### Population allele frequencies/patient cohort data

Patient data, including SOD1-ALS age of onset, disease duration, enzymatic specific activity, and both abundance and half-life (calculated in mice) were obtained from Huang et al. 2024^37^, which aggregated information from the literature. In the case where multiple papers reported data for a given variant (e.g. enzymatic activity of SOD1-L118V was measured as both 61%^38^ and 88%^114^), a weighted average was calculated with weight based on the sample size in each study.

### Structure analyses

We used PyMOL to place map scores in the context of solved crystal structures of apo-SOD1 (PDB ID: 1HL4), holo-SOD1 (PDB ID: 1HL5)^47^, and the Zn^2+^-replete SOD1-CCS complex (PDB ID: 6FP6)^61^. To simulate the binding of the de-metalated intermediate (monomer) of SOD1 (PDB ID: 1RK7)^62^ with CCS (PDB ID: 1DO5)^63^, we used the ClusPro web server (https://cluspro.bu.edu/publications.php) with default parameter settings for protein-protein docking. The monomeric SOD1 and CCS PDB structures were uploaded to the server as the ligand and receptor, respectively. ClusPro initially performs rigid-body docking using the PIPER algorithm, clusters the top 1,000 complexes based on RMSD, then refines and ranks these complexes through energy minimization using the CHARMM force field.

Residues at the interface (within a threshold of 5.0 Å) of homo- and heterodimer complexes were identified using the PRODIGY-crystal (https://wenmr.science.uu.nl/prodigy) web server, which bases its predictions on structural and energetic features ^115,116^. We used the FreeSASA program (https://freesasa.github.io/) to calculate the relative solvent exposure of residue positions. After examining the distribution of relative solvent exposure values, we established thresholds corresponding to the high and low peaks. Residues with surface area values exceeding 35% were considered exposed, while those below 20% were classified as core (“buried”).

### Setup of molecular dynamic simulations

To examine variant-specific impacts on the dynamic behavior of SOD1, especially movements associated with metal-binding, we performed molecular dynamics (MD) simulations using a high-resolution monomeric SOD1 structure (PDB: 2C9V^117^, chain A). The apoenzyme system was constructed by removing metal ions (Zn²⁺ and Cu²⁺) and introducing site-specific mutations with PyMol (version 1.3, Schrödinger, LLC). MD simulations were conducted with the NAMD software suite, employing the CHARMM36m force field, and the system was prepared via the CHARMM webserver (https://www.charmm-gui.org/), reflecting physiological conditions (pH 7.4). The N-terminus and C-terminus were patched to their charged states, and a disulfide bond was established between C58 and C147

The protein was solvated in a rectangular box of TIP3P water molecules with a 10.0 Å minimum distance from the edge of the box. To simulate physiological ionic strength, 0.15 M potassium chloride was added using the Monte Carlo method.

Production simulations were run at 310 K for 400 ns across five independent runs. MD trajectories were analyzed using the Python package ProDy. System stability was assessed by calculating the mean-square fluctuations (MSF) of backbone atoms across the five runs. Secondary structure and flexibility profiles were examined, and hydrogen bond interactions were defined with a 120° angle (donor-hydrogen···acceptor) threshold and a 3.4 Å distance threshold between the donor and acceptor heavy atoms.

## Weblinks

Document S1 (Supplemental Note)

Table S1 (All datasets and primers)

## Data code and availability

Final map scores for both maps, and abundance map LLRp values with confidence intervals and ACMG-compatible evidence strengths are available in Table S1. Final scores are also available on MaveDB^118^ for the both the abundance map including LLRp values and evidence weights (accession number: urn:mavedb:00001217-a-1) and the activity map (accession number: urn:mavedb:00001217-a-2). Code used for analysis can be found on GitHub (https://github.com/axakova/SOD1_Manuscript).

## Acknowledgements

We gratefully acknowledge funding for this project from Biogen Inc. We further acknowledge support from the National Institutes of Health National Human Genome Research Institute (NIH/NHGRI) Center of Excellence in Genomic Science Initiative (<u>HG010461</u>), the NIH/NHGRI Impact of Genomic Variation on Function (IGVF) Initiative (<u>UM1HG011989</u>), and a Canadian Institutes of Health Research Foundation Grant (FDN-159926) to F.P.R. We thank G. Lum and A. Nasrabad for computational assistance related to molecular dynamics simulations.

## Declaration of interests

F.P.R. is an investor in Ranomics, Inc., and is an investor in and advisor for SeqWell, Inc., BioSymetrics, Inc., and Constantiam Biosciences, Inc.

## Notes

https://github.com/axakova/SOD1_Manuscript

https://www.mavedb.org/

### Competing Interest Statement

The authors have declared no competing interest.

