## Supplementary material for "Landscapes of missense variant impact for human superoxide dismutase 1": Document S1

### Supplemental Note

#### Preliminary explorations to find an multiplexed assay of SOD1 gain of function effects

ALS-linked SOD1 variants have been reported to confer a gain of function in that they promote the formation of oligomers[1], with toxicity that correlates with the presence of smaller oligomers rather than larger aggregates[2]. Since SOD1-ALS is a dominant disease, we did not knock out the endogenous *SOD1* in HEK293T cells to better recapitulate the endogenous system. Indeed, one pathomechanism in cultured cells but not patients that has been implicated in ALS is the potential for wild-type SOD1 to heterodimerize with SOD1 variants and thereby increase the amount of aggregated SOD1[3,4]. Although we showed that human cells bearing SOD1-GFP fusions could reliably identify variants that decrease protein abundance, examination of these cells by fluorescence microscopy did not reveal GFP puncta indicating protein aggregation for either gain of function variants or WT cells under the growth conditions we used initially. We therefore sought alternative conditions that might sensitize cells to this toxic aggregation phenotype, with the hope that we might then use FACS to detect aggregating variants[5]. Previous reports suggested that cells can be sensitized to yield SOD1 variant toxicity either by SOD1 variant overexpression, or by environmental changes that either disrupt calcium homeostasis, trigger calpain or activate neuronal nitric oxide synthase[6,7].

First, to assess whether HEK293T cells for could be sensitive to SOD1 aggregation via its overexpression, we used transient transfection of plasmids with SOD1-GFP (WT; pathogenic variants p.Ala5Val, p.Gly86Arg, p.Arg116Gly which are known to aggregate) downstream of the CMV promoter. This was performed both in our baseline growth condition and after addition of the cysteine protease inhibitor MG-101 (also called ALLN) that has been reported to reduce the processing of misfolded SOD1 and result in a higher aggregation[7]. In these experiments, ALLN treatment did yield more aggregation for p.Ala5Val, p.Gly86Arg and p.Arg116Gly variants than for WT SOD1, but aggregation was observed in fewer than 1% of all cells (Figure S3A). The weakness of this effect, coupled with the fact that our multiplexing strategy is most effective when each cell expresses only a single variant, led us to abandon this sensitization strategy.

Second, we assessed whether SOD1 was able to aggregate when more stably expressed in the HEK293T cell system alone, with or without treatment with KNK437, a heat shock protein 70 chaperone inhibitor. HEK293T integrant cells bearing SOD1-GFP for multiple alleles, including WT, known-aggregating variants p.Gly86Arg and p.Leu145Phe, as well as p.Gly130Ser (with an unknown propensity to aggregate). We examined cells for aggregation after 24 and 48 hours. Only for KNK437 (and only after 48 hours) did we observe the aggregating phenotype, and this was observed only in <1% of all cells stably expressing pathogenic variants (Figure S3B). Low penetrance of this cellular phenotype, coupled with variable aggregation across replicates suggested that this assay was unsuitable for a large-scale assay. Additional attempts to increase the sensitivity of this assay to aggregation using combinations of exogenous NO donor (GSNO), calcium ionophore (A23187), sorbitol, heat shock, ALLN and KNK437 were unsuccessful (data not shown).

### Supplemental Figures

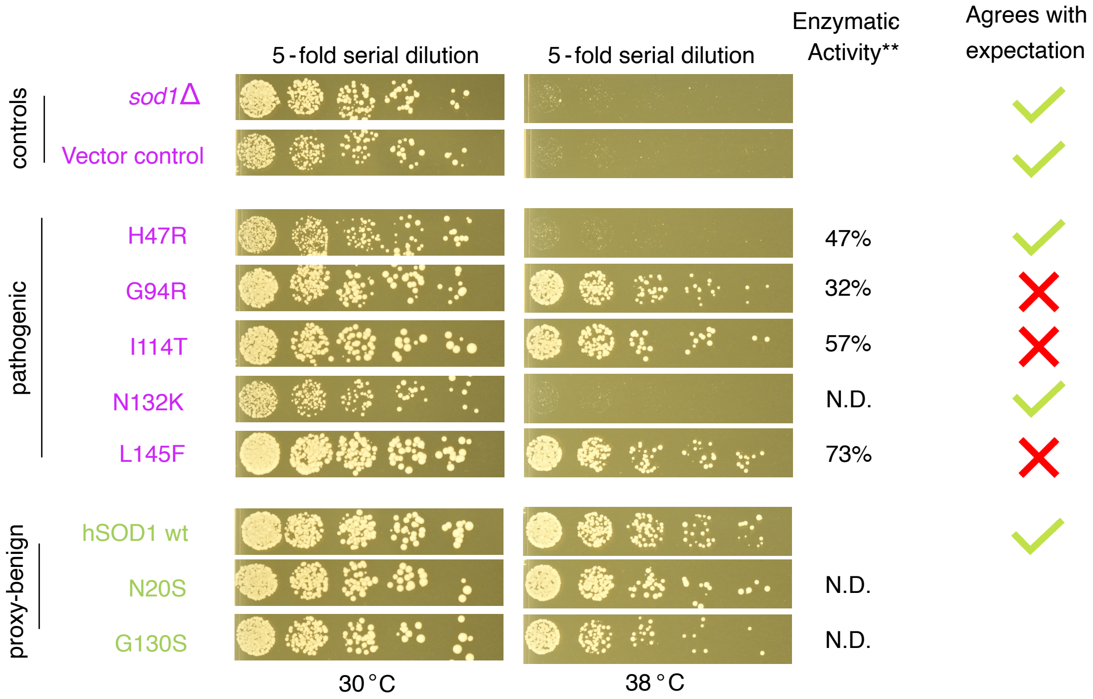


**Figure S1. Functional complementation assay results showing whether expression of human SOD1 protein variants can rescue growth of a yeast *sod1Δ* strain.**
Complementation assay results for ClinVar-reported negative controls and pathogenic variants (purple text), and WT control and proxy-benign variants (green text). Five-fold serial dilutions of yeast cells were spotted onto plates, with growth evaluation after incubating for 48 hours at either permissive (30°C) or non-permissive (38°C) temperature. Patient enzymatic activity measurements[8] for p.His47Arg, p.Gly94Arg, p.Ile114Thr and p.Leu145Phe are shown.

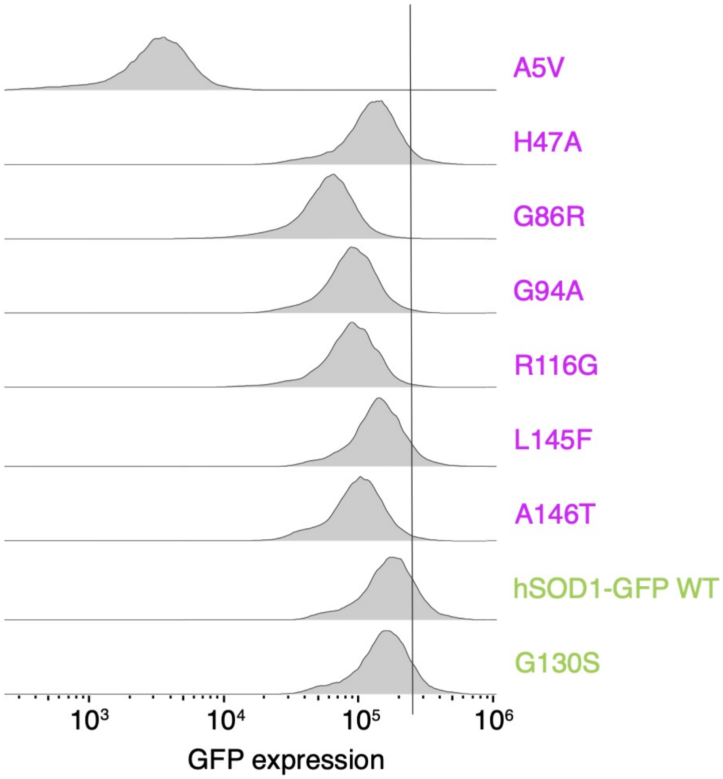


**Figure S2: Distributions of GFP intensity from HEK293T cells with stably integrated SOD1-GFP indicate reduced abundance for pathogenic SOD1 variants.**
Flow cytometric GFP distribution of single, viable cells with integrated SOD1-GFP at the *Bxb1* site in HEK293T cells for pathogenic variants (purple text), and WT SOD1-GFP control and one proxy-benign variant (p.Gly130Ser) taken from gnomAD (green text). Black vertical line indicates potential gating strategy to enrich for variants with high abundance.

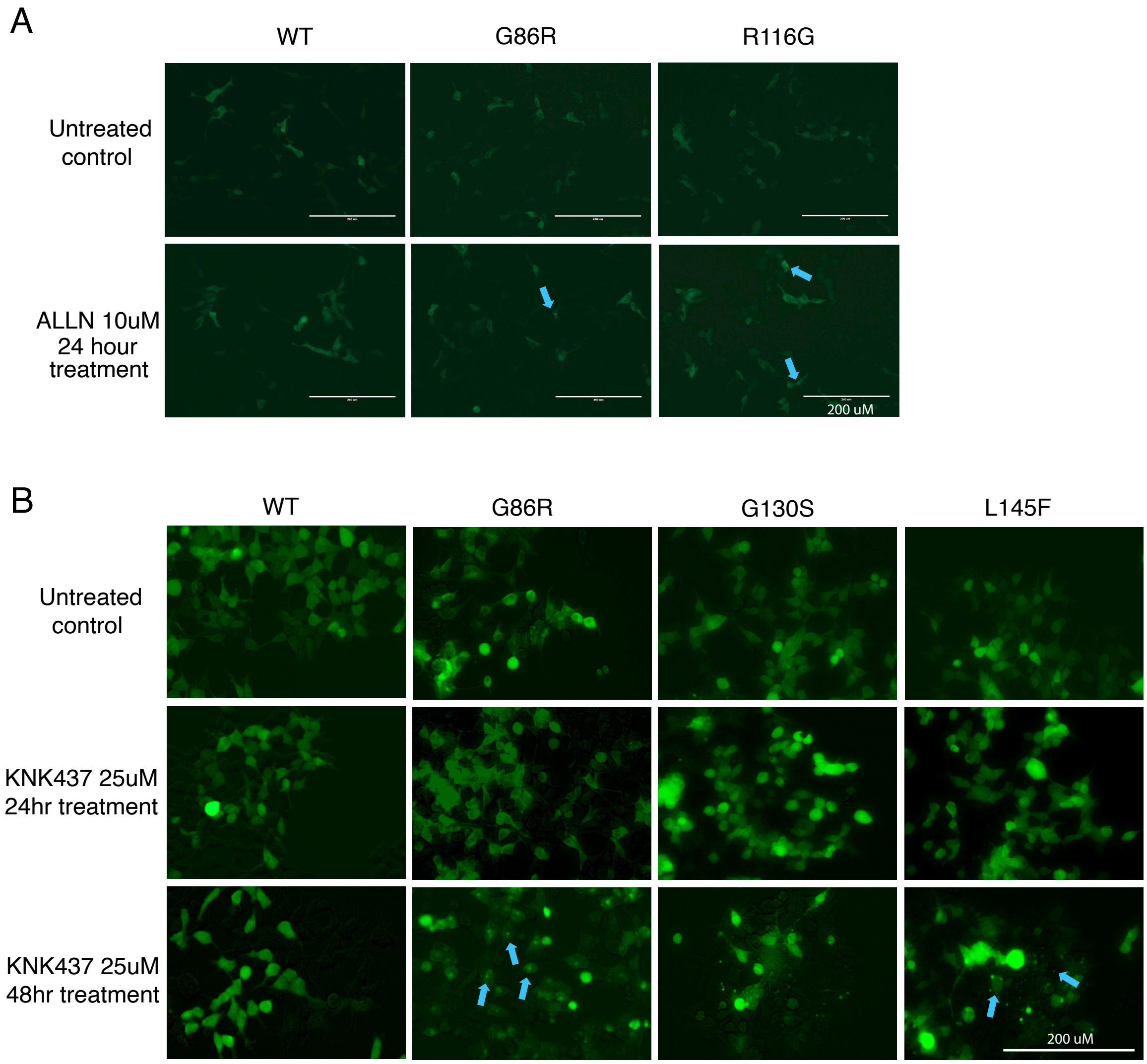


**Figure S3. Chemical treatment of HEK293T cells achieved only limited SOD1-GFP aggregation.** Aggregates indicated by the blue arrows for: (A) HEK293T cells with SOD1-GFP expressed at high levels via transient transfection, with and without ALLN treatment; (B) HEK293T cells with SOD1-GFP expressed at moderate levels via stable integration in the Bxb1 recombination site, with and without KNK437 treatment.

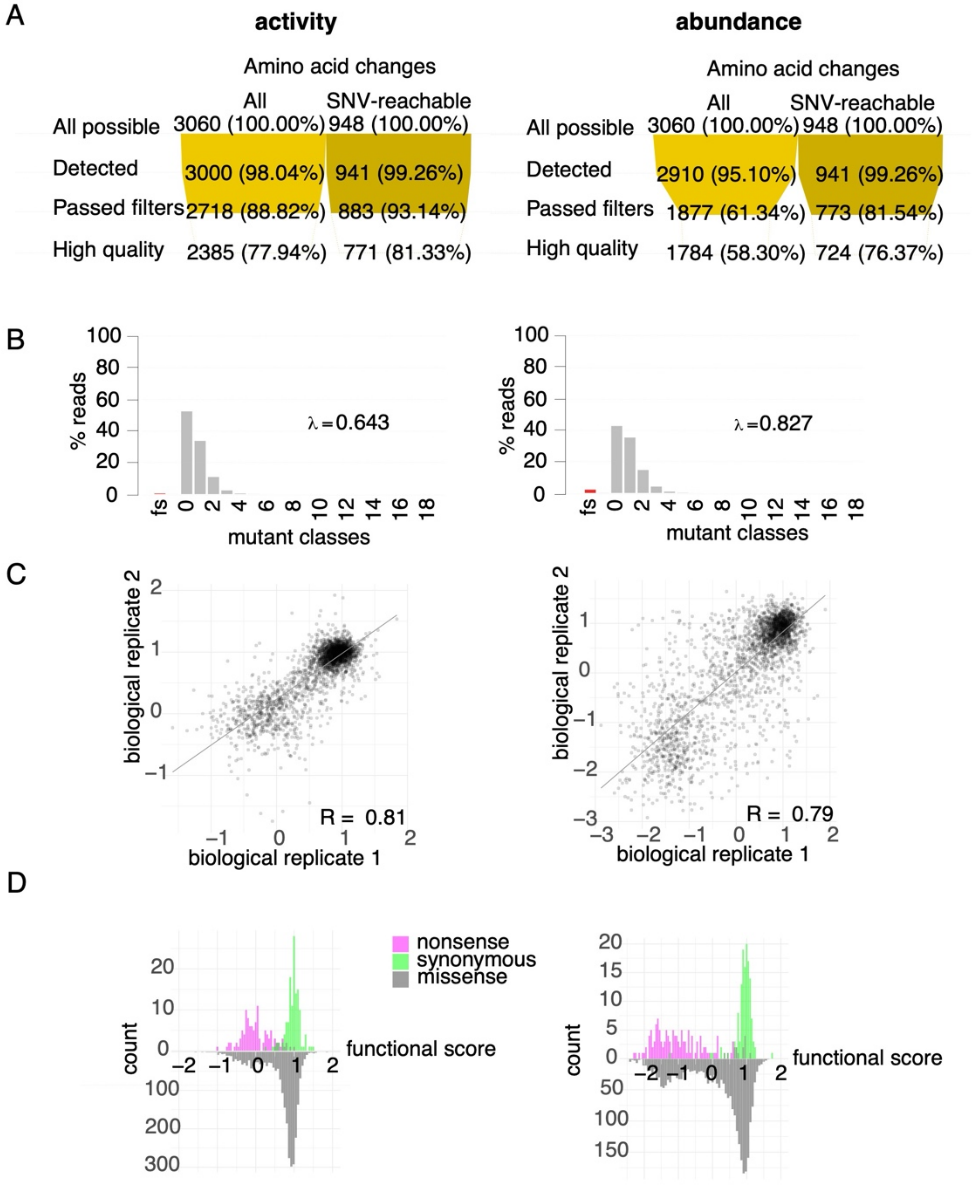


**Figure S4. Characterization of SOD1 variant libraries used for activity (top) and abundance (bottom) variant effect maps.**

(A) Percentage of variants detected and passing quality control are shown for each SOD1 map. For each map, the number of synonymous, nonsense, and missense substitutions are shown across all residue positions, both before (left) and after (right) restricting to substitutions that are possible given a single nucleotide change. The four rows correspond to: 1) theoretically possible substitutions; 2) substitutions detected in the pre-selection condition; 3) substitutions above a threshold pre-selection frequency; and 4) substitutions of high quality (standard error < 0.3).

(B) Distribution of the number of missense variants in clones from the total enzymatic activity and abundance SOD1 mutagenized libraries, and the fraction of clones carrying small indels resulting in frameshifts (“fs”). The average number of amino acid changes per clone (λ) is also estimated (see material and methods).

(C) Biological replicate functional score correlations for each assay.

(D) Distributions of measured functional impact scores for nonsense (pink), synonymous (green), and missense (gray) variants for the maps.

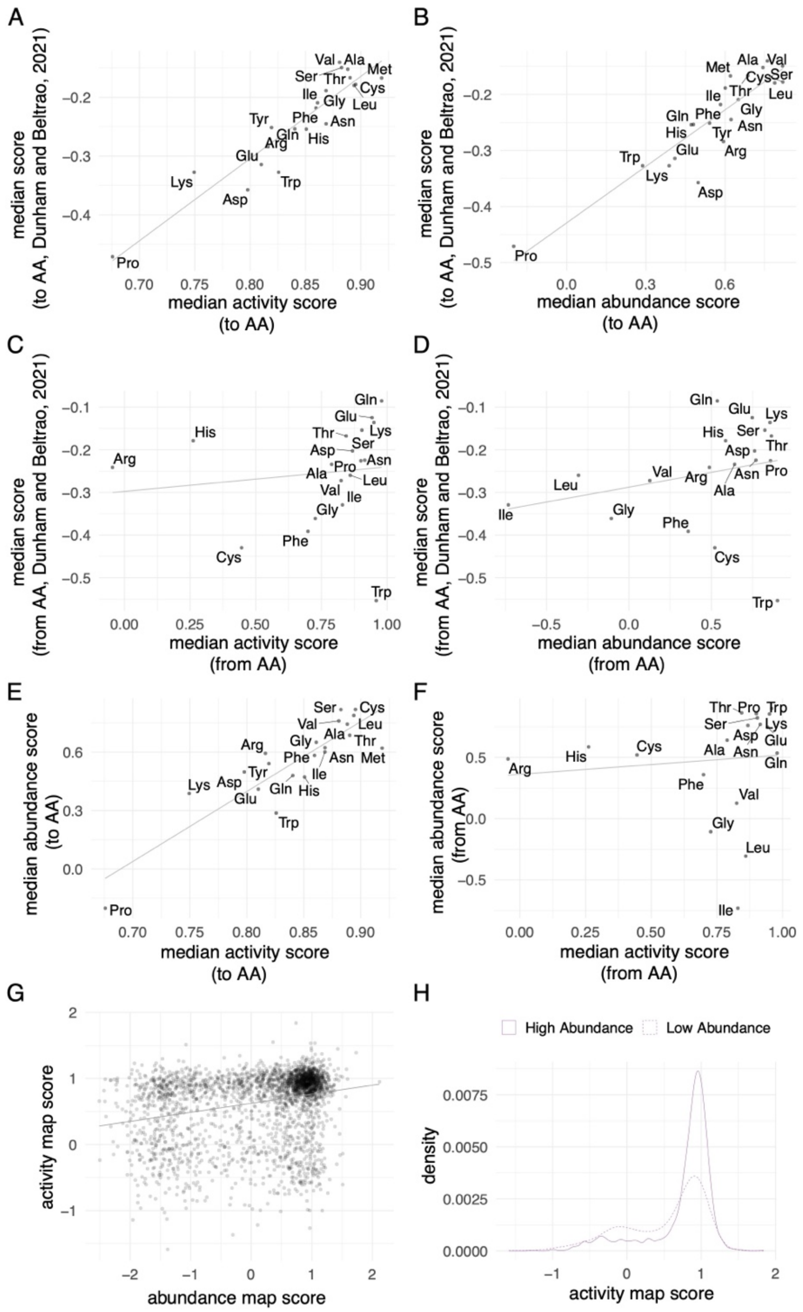


**Figure S5. Correspondence between functional scores of SOD1 missense variants from the total enzymatic activity and abundance maps**.

Correlation of activity and abundance map functional scores to aggregated scores from substitutions to (A-B) and from (C-D) amino acids from 28 deep mutational scans[9]. (To amino acids: Spearman's R activity = 0.89, p<2e-16. Spearman's R abundance = 0.89, p<2e-16. From amino acids: Spearman's R activity = 0.48, p=0.045, abundance p>0.05). Correlation of scores corresponding to median score of changes to (E) and from (F) specific amino acids between abundance and activity map (Spearman's R to amino acid = 0.85, p<2e-16; Spearman's R from amino acid = 0.28, p<2e-16). Correlation between activity and abundance map scores (G; Spearman's R = 0.28; P < 2e-16). Activity scores corresponding to variants with high and low-abundance functional scores (H; abundance score > 0.5, abundance score <= 0.5, respectively.) Solid lines represent linear regression fits.

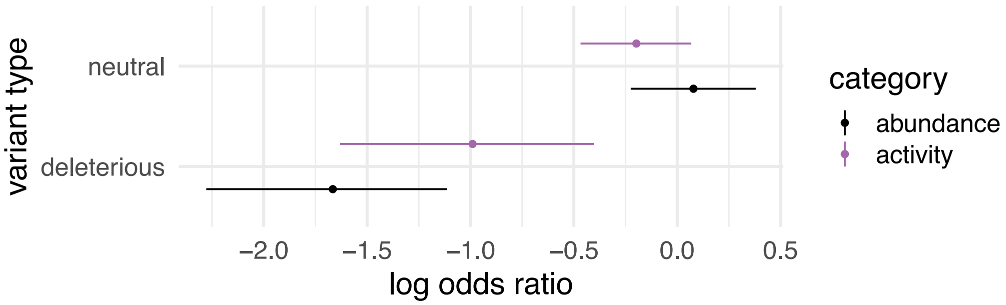


**Figure S6. Depletion in human cohorts of missense variants with damaging functional impact scores for activity and abundance assays.** Log-odds ratios for the depletion of variants with neutral or damaging scores from total enzymatic activity (purple) or abundance (black) maps in both UK Biobank and gnomAD population sequencing databases. Range lines represent 95% confidence intervals.

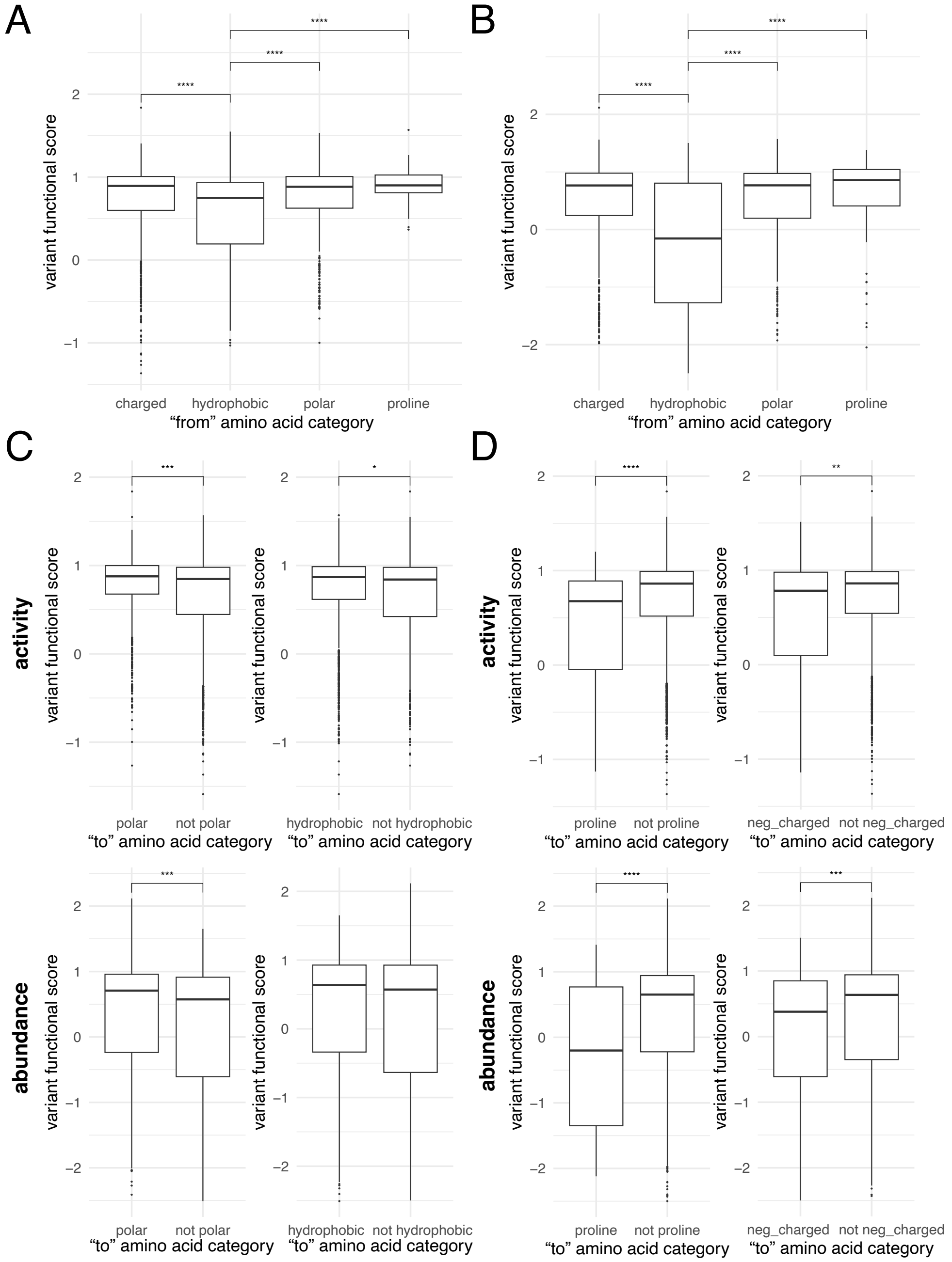


**Figure S7: Variant functional scores by biochemical property.**
Variant functional scores by initial amino acid category for the (A) activity map and (B) the abundance map. Scores for conservative substitutions compared to all else from (C) initially polar positions and initially hydrophobic positions for activity (top) and abundance maps (bottom). Scores for variants changed to proline, or to negatively charged residues (Asp or Glu) for activity (top) and abundance maps (bottom). p**<0.01, p***<0.001, p****<0.0001 by Wilcoxon.

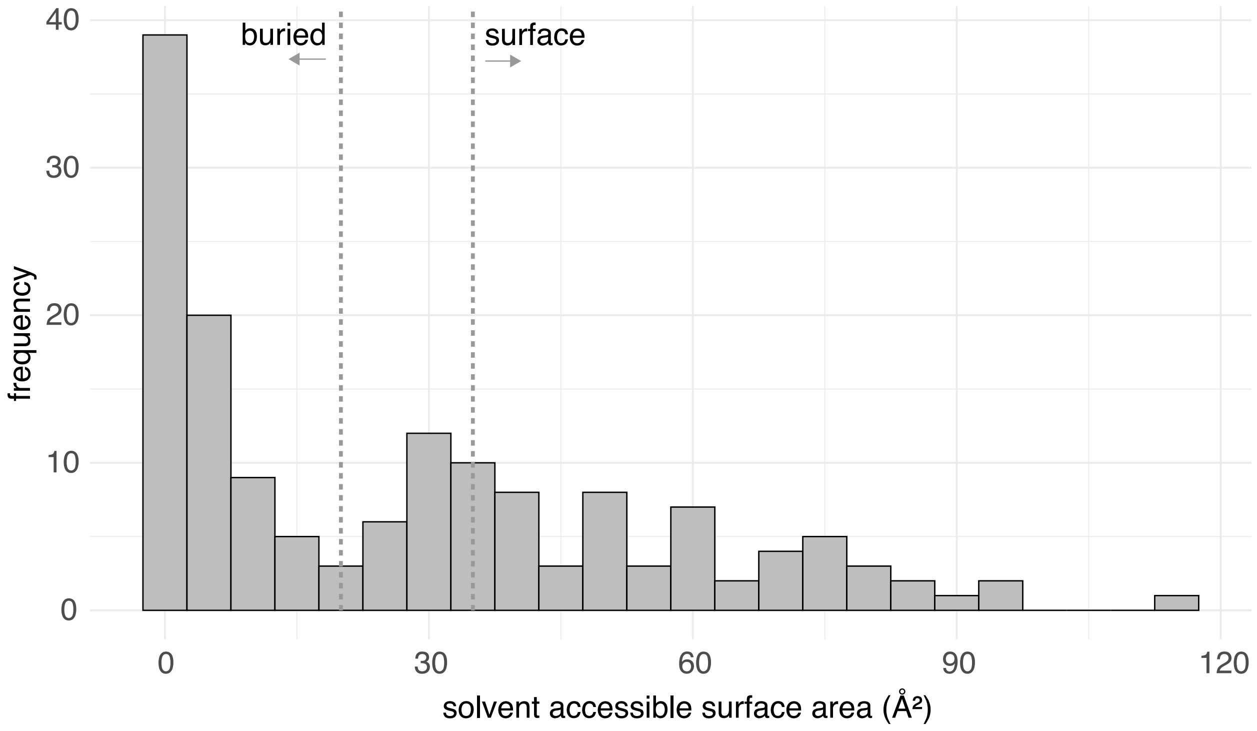


**Figure S8. Distribution of solvent accessible surface area for SOD1 residues.**
SOD1 residues with surface area values exceeding 35% (based on freeSASA values) were considered exposed, while those below 20% were classified as buried (see Methods).

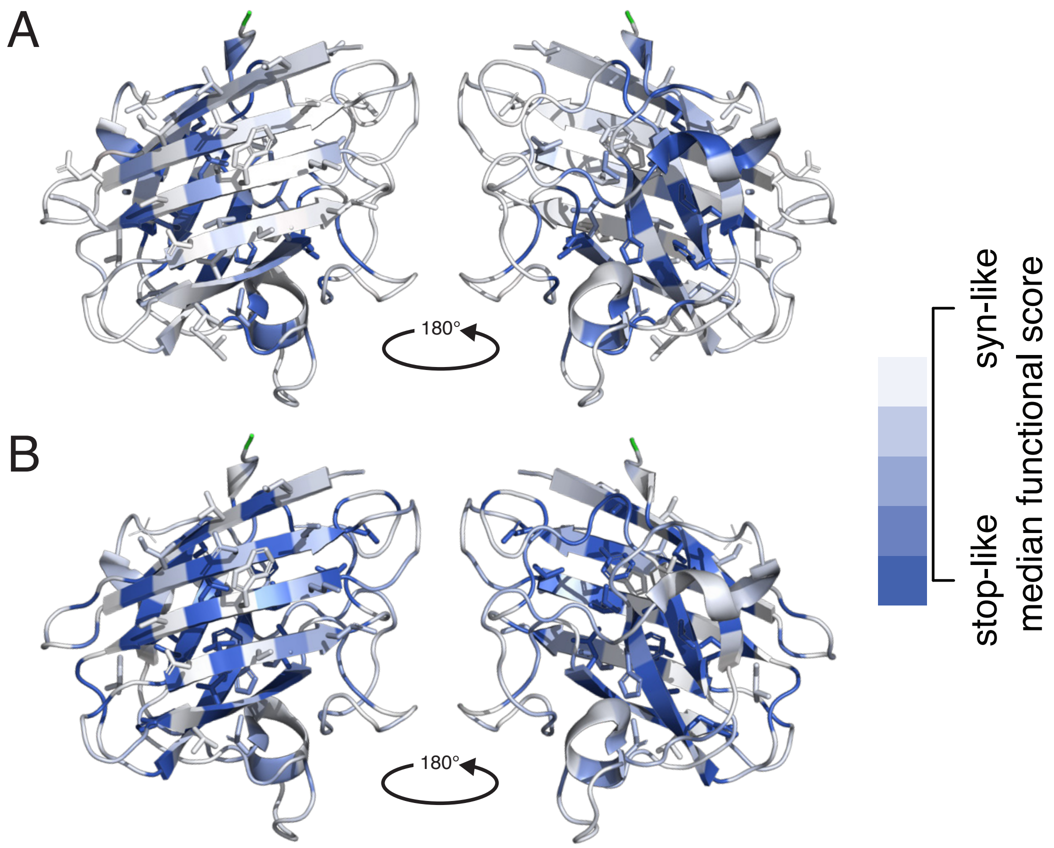


**Figure S9. Crystal structure for SOD1 with residues colored according to map scores.** Residues in the structure, PDB:1HL5 [10], were colored based on the mean score from (A) the enzymatic activity map and (B) the abundance map.

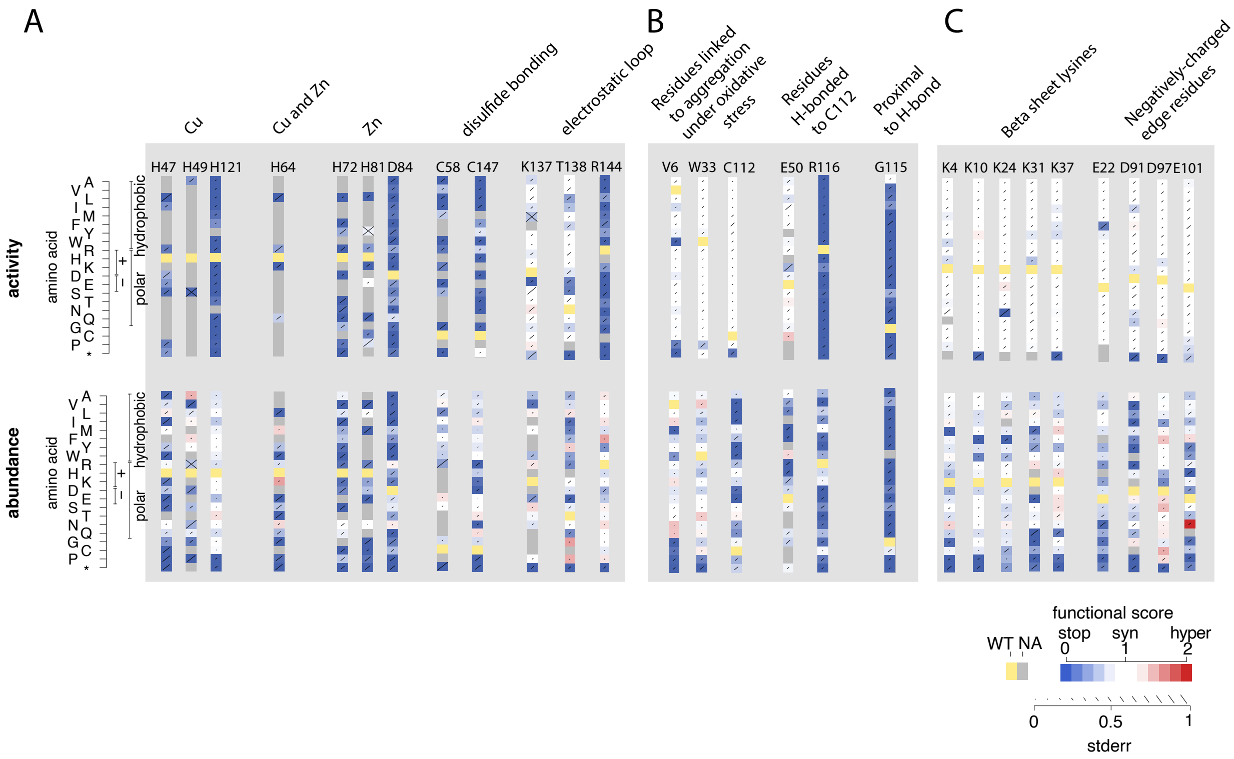


**Figure S10. Identifying patterns of mutational tolerance in SOD1.**
Functional scores for each possible amino acid substitution (y-axis) at specific SOD1 residue position sets of interest (x-axis). Sets of interest included: (A) residues that participate in $Cu^{2+}$ and $Zn^{2+}$-binding, disulphide bonds, or the electrostatic loop that promotes high-affinity site-specific metal binding; (B) residues at which variation has been reported to either cause SOD1 aggregation under oxidative stress, or instability of the H-bond around p.Cys112; and (C) lysine residues within the first β-sheet and negatively-charged residues at the edge of the first β-sheet.

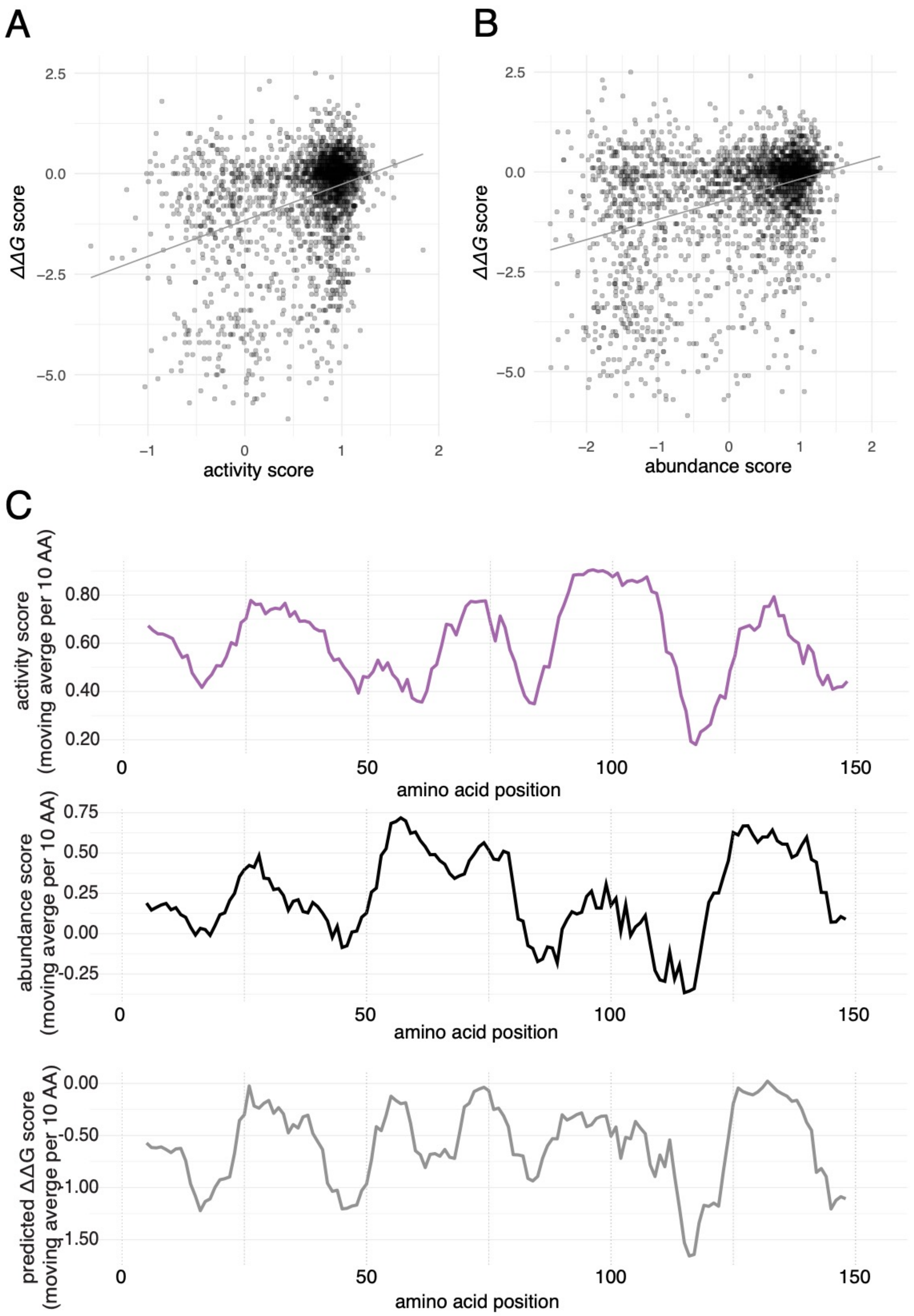


**Figure S11. Comparing map scores with predicted effects on protein stability.** Functional scores compared to free energy change (*ΔΔG*) for activity (A) and abundance (B) maps. (C) Moving windows of scores from the total enzymatic activity (purple), abundance maps (black), and predicted *ΔΔG* (light grey) values for SOD1 missense variants at different protein positions. Plotted values represent averages within windows of ten amino acid (AA) positions.

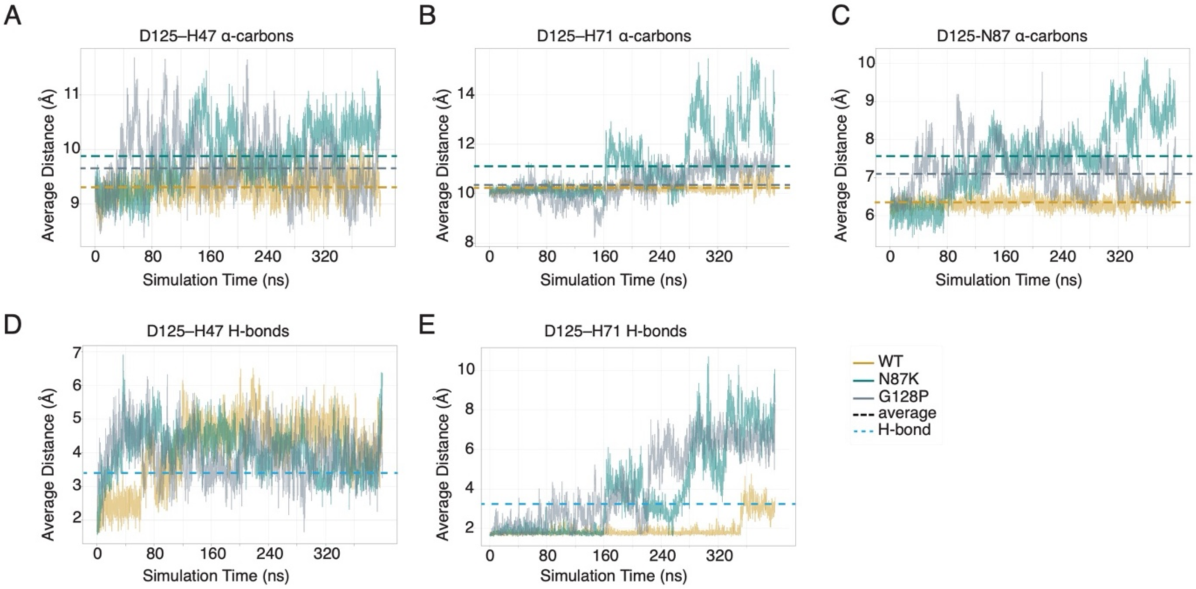


**Figure S12. Molecular dynamics simulations evaluating impact of variation on inter-residue distances relevant for SOD1's ability to bind metal.** The dotted lines represent the average distances between the α-carbons of residue 125 and (A) p.His47, (B) p.His72, and (C) p.Asn87 in WT SOD1, as well as in the p.Asn87Lys and p.Gly128Pro variants. The blue dotted line indicates the hydrogen bond interaction threshold (3.4 Å) for p.Asp125-p.His47 and p.Asp125-p.His72.

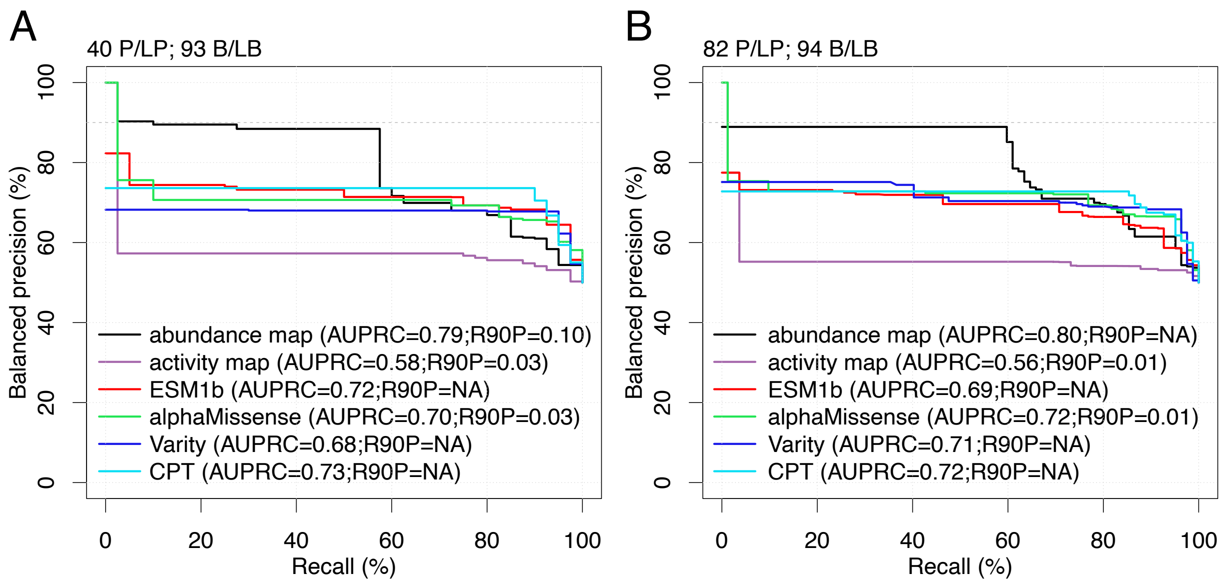


**Figure S13. Comparison of total enzymatic activity and abundance SOD1 maps with computational predictors using balanced precision recall analysis for SOD1**. We used positive reference variants from (A) ClinVar and (B) Labcorp Genetics databases, and negative reference variants from gnomAD to draw PRC curves for the maps and computational predictors. Here we evaluate precision (fraction of variants scoring below each threshold functional impact score that are in the positive reference set containing pathogenic variants) vs recall (fraction of positive reference variants with functional scores below threshold). Precision has been transformed to reflect performance in a balanced test setting where positive and negative sets contain the same number of variants. Balanced precision-recall curves are shown for the total enzymatic activity (purple) and abundance maps (black), as well as computational predictors; ESM1b (red), AlphaMissense (green), VARITY (blue) and CPT (turquoise). Positive and negative reference set sizes (P/LP and B/LB, respectively; see Methods) are indicated. The number of variants in the reference set included in these PRC curves was lower than the curated reference sets due to the intersection between sets for experimental and computational approaches. The variant p.Asn20Ser was classified as benign by Labcorp Genetics but was excluded from the ClinVar set because of a conflicting annotation.

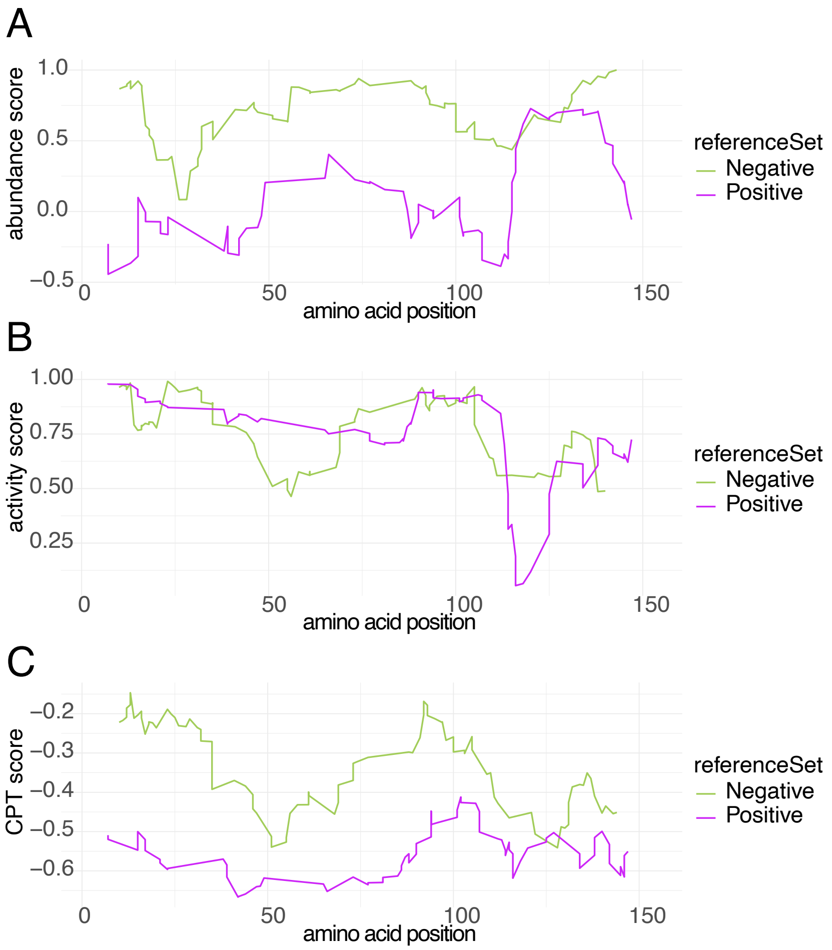


**Figure S14**. **Correspondence of SOD1 maps and computational predictor CPT to clinical variant annotations.** Plotted values are (A) abundance map, (B) activity map and (C) computational predictor CPT scores as part of the positive (magenta) or negative (green) reference variant sets from Labcorp Genetics or gnomAD proxy benign, repectively.

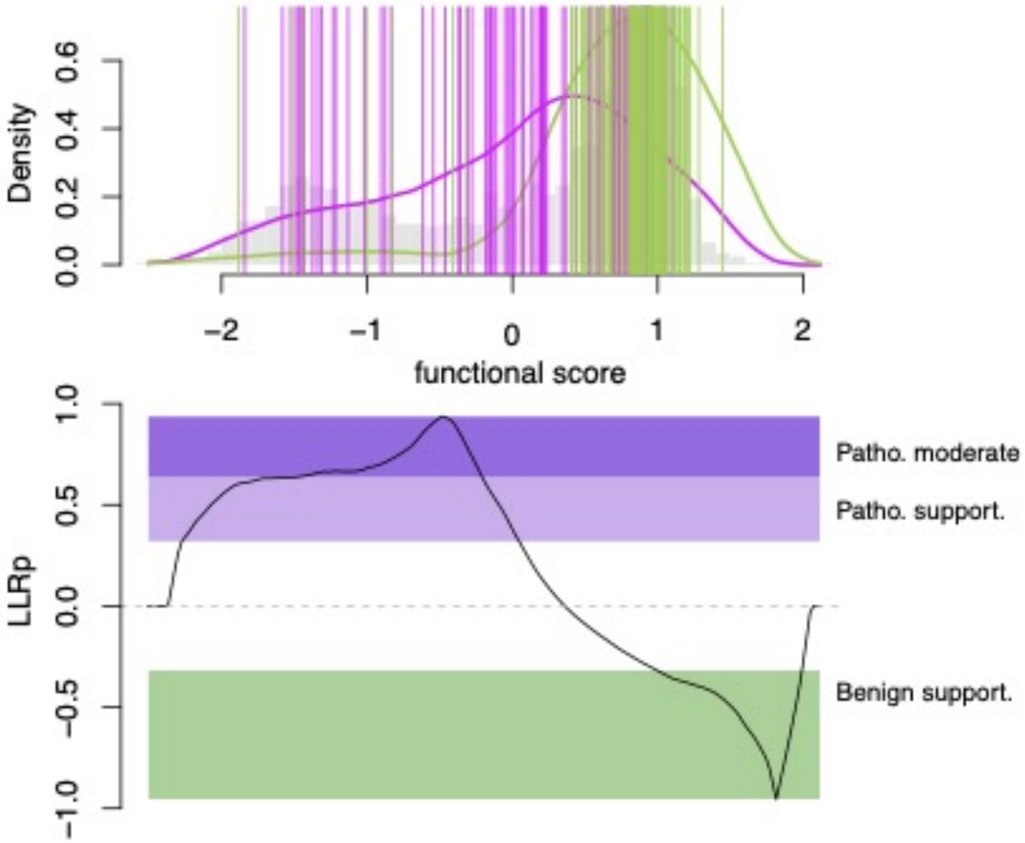


**Figure S15. Evidentiary value of abundance map scores for clinical variant interpretation.** Calculation and distribution of log-likelihood ratios (LLR) of pathogenicity for scores of the SOD1 abundance map. The functions (top) express the log ratio between the likelihood of observing a given score in the score distribution of the positive reference set (magenta) from Labcorp Genetics as opposed to that of the negative reference set (green). Gray histogram bars show the distribution of missense variants for comparison.

**Supplemental Table 1:** Summary table of molecular dynamics simulations for residues involved in SOD1 electrostatic loop.

| | | | Between Position 1 and 2 | |
|---|---|---|---|---|
| **Condition** | **Position 1** | **Position 2** | **Ca Distance (A)** | **Hydrogen Bonding Time (%)** |
| WT | p.Asn87 | p.Asp125 | 6.4 ± 0.2 | |
| WT | p.His47 | p.Asp125 | 9.3 ± 0.3 | 22 |
| WT | p.His72 | p.Asp125 | 10.2 ± 0.2 | 98 |
| p.Asn87Lys | p.Asn87Lys | p.Asp125 | 7.6 ± 1.0 | |
| p.Asn87Lys | p.His47 | p.Asp125 | 9.9 ± 0.6 | 18 |
| p.Asn87Lys | p.His72 | p.Asp125 | 11.2 ± 1.4 | 53 |
| p.Gly128Pro | p.Asn87 | p.Asp125 | 7.1 ± 0.5 | |
| p.Gly128Pro | p.His47 | p.Asp125 | 9.7 ± 0.5 | 22 |
| p.Gly128Pro | p.His72 | p.Asp125 | 10.4 ± 0.6 | 46 |
